# Developmentally programmed nuclear pore complex replacement enables oocyte specification

**DOI:** 10.64898/2026.02.13.705775

**Authors:** Shruti Venkat, Tram Nguyen, Cecilia Blangini, Michelle Pollak, Karen Schindler, Maya Capelson, Prashanth Rangan

**Affiliations:** Department of Stem Cell Biology and Regenerative Medicine, Black Family Stem Cell Institute, Icahn School of Medicine at Mount Sinai, New York, NY 10029, USA; Cell and Molecular Biology Program, Department of Biology, San Diego State University, San Diego, CA 92182, USA; Department of Genetics, Rutgers, The State University of New Jersey, Piscataway, NJ 08854

**Keywords:** Nuclear pore complex, ESCRT-III, Vps4, oocyte specification, Drosophila, oogenesis, nuclear remodeling, maternal inheritance, cell fate transition

## Abstract

Oocytes endow embryos with molecular machinery essential for development, but not all maternal components are inherited indiscriminately. In Drosophila, surveillance pathways eliminate defective mitochondria and aberrant RNAs from the maternal pool. Whether stable nuclear structures, like nuclear pore complexes (NPCs), are similarly curated remains unknown. Here, we uncover a developmentally programmed NPC turnover pathway that renews NPCs during oocyte specification. NPC levels decline through a combination of passive dilution, driven by deferred nucleoporin expression, and active degradation mediated by the ESCRT-III/Vps4 pathway. This clearance is counterbalanced by subsequent de novo NPC synthesis. Failure to turn over NPCs results in aberrantly persistent germ cell gene expression and defective oocyte specification. These findings establish NPC renewal as a critical step in oocyte identity establishment and maternal provisioning.

## Introduction

Every generation begins with a renewed cellular state that supports faithful development. This renewal depends on newly synthesized maternal components deposited into the egg, including mitochondria, RNAs, proteins, and large macromolecular assemblies that sustain early embryogenesis prior to zygotic transcription (*1*, *2*). Increasing evidence indicates that maternal contribution is actively curated, as oocytes selectively eliminate damaged mitochondria and inappropriate transcripts to establish developmental competence (*3–5*). Whether other essential macromolecular structures, such as nuclear pore complexes (NPCs), are similarly regulated during oogenesis remains unclear.

NPCs are large protein assemblies embedded in the nuclear envelope that regulate nucleocytoplasmic transport and chromatin organization to influence gene expression (*6*, *7*). Each NPC is composed of multiple nucleoporins (Nups) organized into distinct subcomplexes, including a channel-forming scaffold and associated structures, that confer both transport function and structural integrity (**Fig. 1A**) (*8*). Many Nups of the core NPC scaffold are exceptionally stable and can persist in post-mitotic cells for years (*9*, *10*). Although this stability is essential for NPC function, it also renders the complex vulnerable to cumulative damage, particularly in long-lived cells such as neurons and oocytes (*11*). In yeast, selective NPC clearance depends on the ESCRT-III/Vps4 pathway, and studies in metazoan disease models and cultured somatic cells suggest that NPC turnover can occur through conserved mechanisms (*12*, *13*). Whether oocytes, which must transmit a functionally intact nuclear environment to the next generation, remodel or renew their nuclear compartment during development remains unknown.

**Figure 1.**
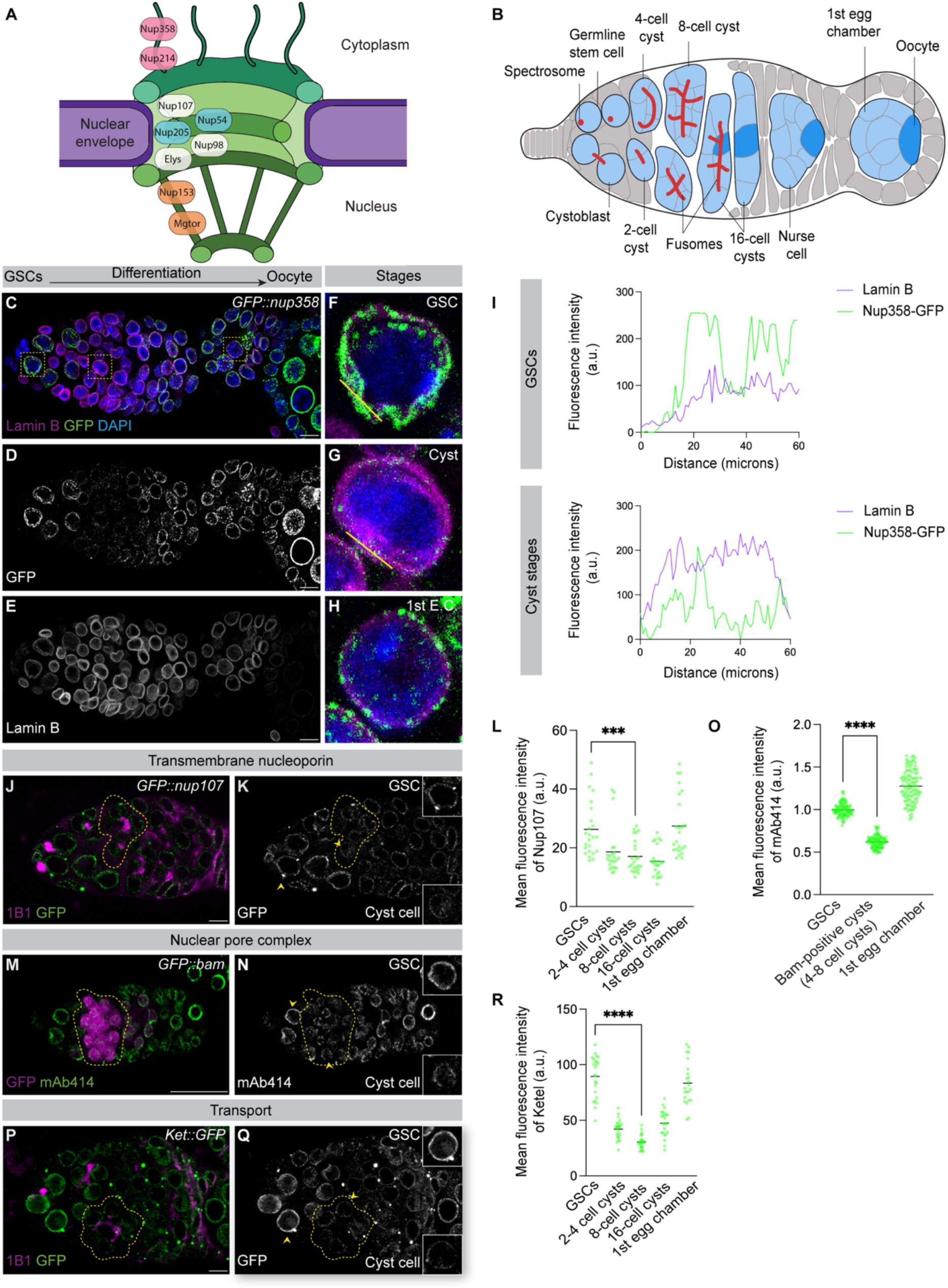
NPC and nucleoporin marker signal is reduced during cyst differentiation preceding oocyte specification. **(A)** Schematic representation of a nuclear pore complex (NPC). **(B)** Schematic of the *Drosophila* germarium. **(C–E)** Confocal images of *GFP::nup358* in the germarium. Scale bar, 10 µm. **(F–H)** Enlarged insets from boxed regions in (C–E), highlighting the nuclear envelope–associated GFP::Nup358 signal in germline stem cells and differentiating cysts. Scale bar, 10 µm. **(I)** Line-scan quantification of Lamin B and GFP::Nup358 fluorescence intensity across nuclei indicated in (F–G). **(J–K)** Confocal images of *nup107::GFP* germaria; gray channel shows Nup107 signal, with dotted yellow line indicating reduced signal. Scale bar, 5 µm. **(L)** Quantification of mean Nup107 fluorescence intensity (a.u.); *n* = 5 germaria; **** = *p* = 0.0002. **(M–N)** Confocal images of *GFP::bam* germaria with mAb414 staining (gray channel) marking NPCs; dotted yellow line indicates regions of reduced mAb414 signal. Scale bar, 25 µm. **(O)** Quantification of mean mAb414 fluorescence intensity (arbitrary units, a.u.). **(P–Q)** Confocal images of *Ket::GFP* germaria; gray channel shows Ket signal, with dotted yellow line indicating reduced fluorescence. Scale bar, 5 µm. **(R)** Quantification of mean Ket fluorescence intensity (a.u.). Statistical analysis where applicable was performed using a two-tailed unpaired Welch’s *t*-test on mean fluorescence intensity. *n* = 5 germaria; **** = *p* < 0.0001 unless otherwise indicated.

Here, we use *Drosophila* oogenesis to investigate whether NPCs are developmentally remodeled during maternal inheritance. In the germarium, germline stem cells (GSCs) divide asymmetrically to produce cystoblasts, which undergo four rounds of incomplete mitotic divisions to form 16-cell cysts (**Fig. 1B**) (*14*). Oocyte specification initiates around the 8-cell stage and is accompanied by chromatin reorganization and transcriptional silencing of germ cell genes (*15*, *16*). By the 16-cell stage, one cell adopts oocyte identity while the remaining cells differentiate as nurse cells (**Fig. 1B**) (*17*). The oocyte remains largely transcriptionally quiescent and depends on maternally deposited components synthesized by nurse cells (*1*, *17*). Although nurse cells are known to synthesize new NPCs and transfer them into the oocyte (*18*), it is unclear whether NPCs assembled during earlier germ cell stages persist. Here, we demonstrate that these pre-existing NPCs are not retained but instead undergo developmentally programmed turnover that is essential for oocyte specification.

## Results

### NPCs accumulate in the oocyte and are predominantly maternally inherited

To determine whether NPCs are maternally inherited, we examined NPC dynamics during oogenesis using complementary genetic and biochemical markers. We visualized distinct NPC substructures using GFP-tagged Nups including Nup358 (cytoplasmic filaments) and Nup107 (core scaffold), together with the mAb414 antibody to detect Nups of the central transport channel that contain Phenylalanine-Glycine (FG) repeats and fluorescent Wheat Germ Agglutinin (WGA) to label glycosylated Nups within the nuclear basket and central channel (*18*, *19*). Using these markers, we observed robust NPC localization at the nuclear envelopes of oocytes, nurse cells, and surrounding somatic follicle cells. NPC components also accumulated within the ooplasm of developing oocytes, consistent with their storage in annulate lamellae for maternal deposition (**Fig. 1S1A-E**) (*18*).

To assess paternal inheritance, we examined spermatogenesis using GFP-tagged Nup107 and Nup358. NPCs were readily detected in germline stem cells and early spermatogenic stages but were largely absent from elongating and mature spermatids at the posterior end of the testis (**Fig. 1S1F–J**). Consistent with this, mAb414-positive NPCs were not readily detected in mature sperm marked by Don Juan::GFP in the seminal vesicle (**Fig. 1S1K–M**) (*20*). Together, these data indicate that NPCs are predominantly maternally inherited, consistent with prior observations (*18*).

### NPC abundance transiently declines during cyst differentiation

Mitochondria and maternally deposited mRNAs are dynamically regulated during cyst stages of oogenesis (*3*, *4*). We therefore asked whether maternally inherited NPCs are similarly regulated during oocyte specification. To exclude effects of mitosis, we assessed nuclear integrity by co-staining germ cells for Lamin B concurrently with the NPC marker GFP::Nup358 (*18*, *21*). Lamin B levels remained stable throughout cyst stages, whereas Nup358 signal intensity at the nuclear envelope decreased markedly (**Fig. 1C–I**), indicating a specific reduction in Nup358 abundance during cyst differentiation. This decline occurred in the absence of phospho-histone H3–positive cells (**Fig. 1S2A–B**), demonstrating that Nup358 loss is not temporally correlated with mitosis.

To determine whether this reduction was specific to Nup358 or reflected broader regulation of the NPCs, we examined Nups representing distinct NPC subcomplexes. In addition to the cytoplasmic Nup358, we analyzed the outer ring components Nup107 and Elys and the nuclear basket Nup153 using a combination of endogenously tagged alleles and antibodies together with the fusome marker, 1B1, to identify the cyst stages (*22*, *23*). All Nups tested exhibited a similar decline during cyst differentiation, reaching a minimum around the 8-cell stage, coincident with the onset of oocyte specification (**Fig. 1J–L**, **Fig. 1S2C–K**). To further confirm that this coordinated reduction reflects decreased NPC abundance rather than selective loss of individual components, we examined pan-NPC markers, including WGA and mAb414, together with a Bag of marbles (Bam::GFP) reporter to stage cysts preceding oocyte specification (*24*). Pan-NPC marker signal was lowest in Bam-positive cyst stages and increased again in later egg chambers (**Fig. 1M–O**, **Fig. 1S2L–N)**, revealing a transient, stage-specific reduction in NPC abundance at the nuclear envelope preceding oocyte specification.

To determine whether NPC reduction is spatially restricted to the future oocyte or reflects a cyst-wide program, we co-stained germaria for WGA and the oocyte marker Orb (*25*). NPC marker signal was comparable between the Orb-positive oocyte and surrounding nurse cells within 16-cell cysts (**Fig. 1S2O–Q**), indicating that NPC reduction occurs uniformly across the cyst. In parallel with reduced NPC marker signal, the nuclear import receptors Ketel (Ket) and Transportin-SR (Tnpo-SR) were also reduced during cyst stages (**Fig. 1P–R**, **Fig. 1S2R–T**) (*26*, *27*). Together, these findings indicate that cyst differentiation is marked by a coordinated, transient reduction in NPC abundance and nuclear transport capacity during oocyte specification.

### Differentiating cyst cells lose pre-existing NPCs and incorporate newly synthesized NPCs

To determine whether NPC reduction is developmentally programmed rather than time-dependent, we analyzed NPC marker signal in germ cells arrested at defined stages of oogenesis. GSCs are long-lived and self-renewing, whereas differentiating cysts undergo defined transitions in developmental identity. Depletion of *bam*, which blocks differentiation and leads to the accumulation of undifferentiated cystoblasts (*24*), resulted in cells that retained NPC marker signal comparable to GSCs (**Fig. 2A–F, J**), indicating that NPC reduction does not occur in undifferentiated germ cells. In contrast, depletion of *RNA-binding Fox protein 1* (*rbfox1*), which is required for the transition from 8- to 16-cell cysts (*28*), led to the accumulation of 8-cell cysts with reduced NPC marker signal (**Fig. 2G–J**). Together, these findings indicate that NPC reduction is initiated during cyst divisions and is coupled to germ cell differentiation rather than chronological time.

**Figure 2.**
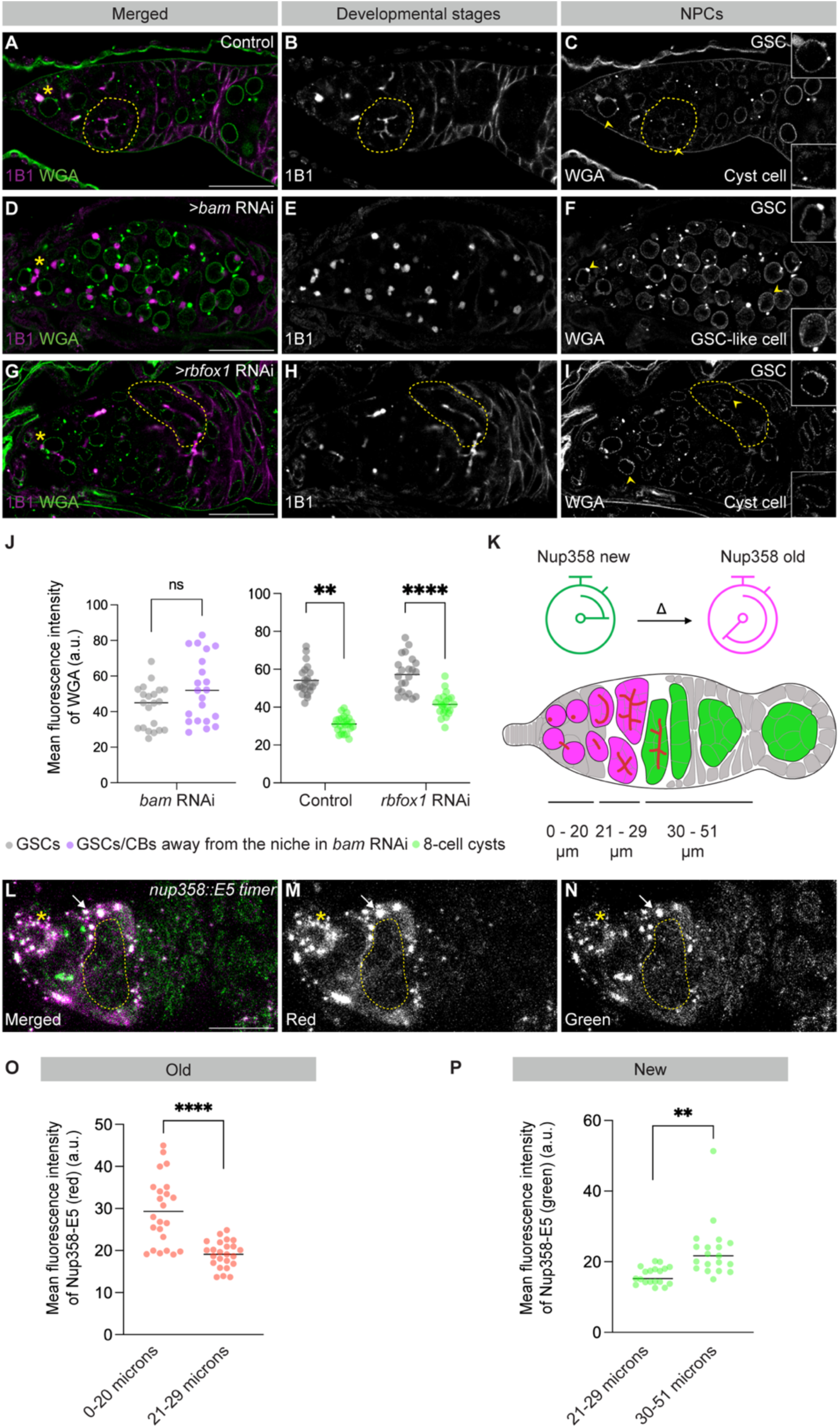
Pre-existing NPCs are selectively lost and replaced with newly synthesized NPCs during cyst differentiation. **(A–I)** Confocal images of control, *bam* RNAi, and *rbfox1* RNAi germaria stained with WGA and 1B1. *bam* RNAi ovaries are enriched for cystoblasts, whereas *rbfox1* RNAi ovaries are enriched for 8-cell cyst stages. Grayscale channels of WGA in *bam* RNAi germaria show maintained NPC marker signal in the absence of cyst differentiation (D), whereas WGA in *rbfox1* RNAi germaria shows reduced NPC marker signal in 8-cell cyst cyst stages relative to GSCs (I). Scale bars, 25 µm. **(J)** Quantification of mean WGA fluorescence intensity. Statistical analysis was performed using a two-tailed unpaired Welch’s *t-*test. *n* = 5 germaria; **** = *p* < 0.0001; ** = *p* = 0.0039. **(K)** Schematic of the E5 fluorescent timer strategy and distance ranges used to assign germline stages. **(L)** Confocal image of a *nup358::E5* germarium showing endogenous E5 fluorescence. Scale bar, 25 µm. **(M)** Grayscale image of the red channel, marking older *nup358::E5* signal enriched in GSCs and early cysts. **(N)** Grayscale image of the green channel, marking newly synthesized *nup358::E5* signal enriched in later cyst stages. **(O–P)** Quantification of mean red and green Nup358::E5 fluorescence intensity across germline stages (arbitrary units, a.u.). Statistical analysis was performed using a two-tailed unpaired Welch’s *t*-test. *n* = 5 germaria; **** = *p* < 0.0001; ** = *p* = 0.0012. Yellow dotted lines indicate cyst stages. Yellow asterisks mark GSCs.

Having established that NPC reduction is developmentally programmed, we next asked whether this decrease reflects preferential loss of NPCs assembled during earlier germ cell stages. To address this, we generated a fly line in which the endogenous *nup358* locus was tagged with the E5 fluorescent timer, which shifts from green to red fluorescence as proteins persist over time (**Fig. 2K**) (*29*). Newly synthesized proteins fluoresce green, whereas older proteins fluoresce red; continuous synthesis yields yellow signal. Homozygous *nup358::E5* flies were viable, indicating that the tag did not disrupt protein function. In somatic cells and GSCs, Nup358::E5 fluorescence was predominantly yellow, consistent with a mixture of newly synthesized and older NPCs (**Fig. 2L-N**). During cyst stages, red fluorescence diminished, indicating loss of NPCs assembled during earlier stages (**Fig. 2M, O**). In early egg chambers, green fluorescence predominated, consistent with incorporation of newly synthesized Nup358 into newly assembled NPCs (**Fig. 2N, P**). Together, these findings indicate that NPC reduction during cyst differentiation involves selective loss of pre-existing NPCs followed by incorporation of newly synthesized components at the onset of oocyte specification.

### Nucleoporin transcription and translation are developmentally delayed during cyst differentiation

The reduction of NPCs assembled during earlier germ cell stages could result from dilution, active removal, or both. If cyst divisions occur without concurrent NPC synthesis, NPCs would be diluted with each division. This model predicts delayed Nup transcription and translation prior to oocyte specification, followed by renewed synthesis at later stages. To test this, we examined the timing of Nup transcription using a validated dual-color transcriptional timer (TransTimer) under the control of GAL4 driven by the endogenous *nup54* promoter (**Fig. 3A**) (*30*). This TransTimer combines a short-lived destabilized GFP and a long-lived nuclear RFP (**Fig. 3A**) (*30*). Somatic cells displayed yellow fluorescence, indicating ongoing transcription, whereas GSCs and early cysts showed attenuated signal (**Fig. 3B–E**). Green fluorescence emerged in 4-cell cysts, and robust green and red fluorescence was detected in 8-cell cysts, indicating strong and sustained *nup54* transcription during cyst differentiation, consistent with our previous finding that heterochromatin formation is required for this upregulation (*15*). Although fluorescence was low in GSCs, *nup54*GAL4-driven depletion of *bam* resulted in accumulation of cystoblasts, consistent with low-level *nup54* transcription in undifferentiated germ cells (**Fig. 3S1A–B**). Together, these data indicate that *nup54* transcription is developmentally deferred and activated during cyst stages.

**Figure 3:**
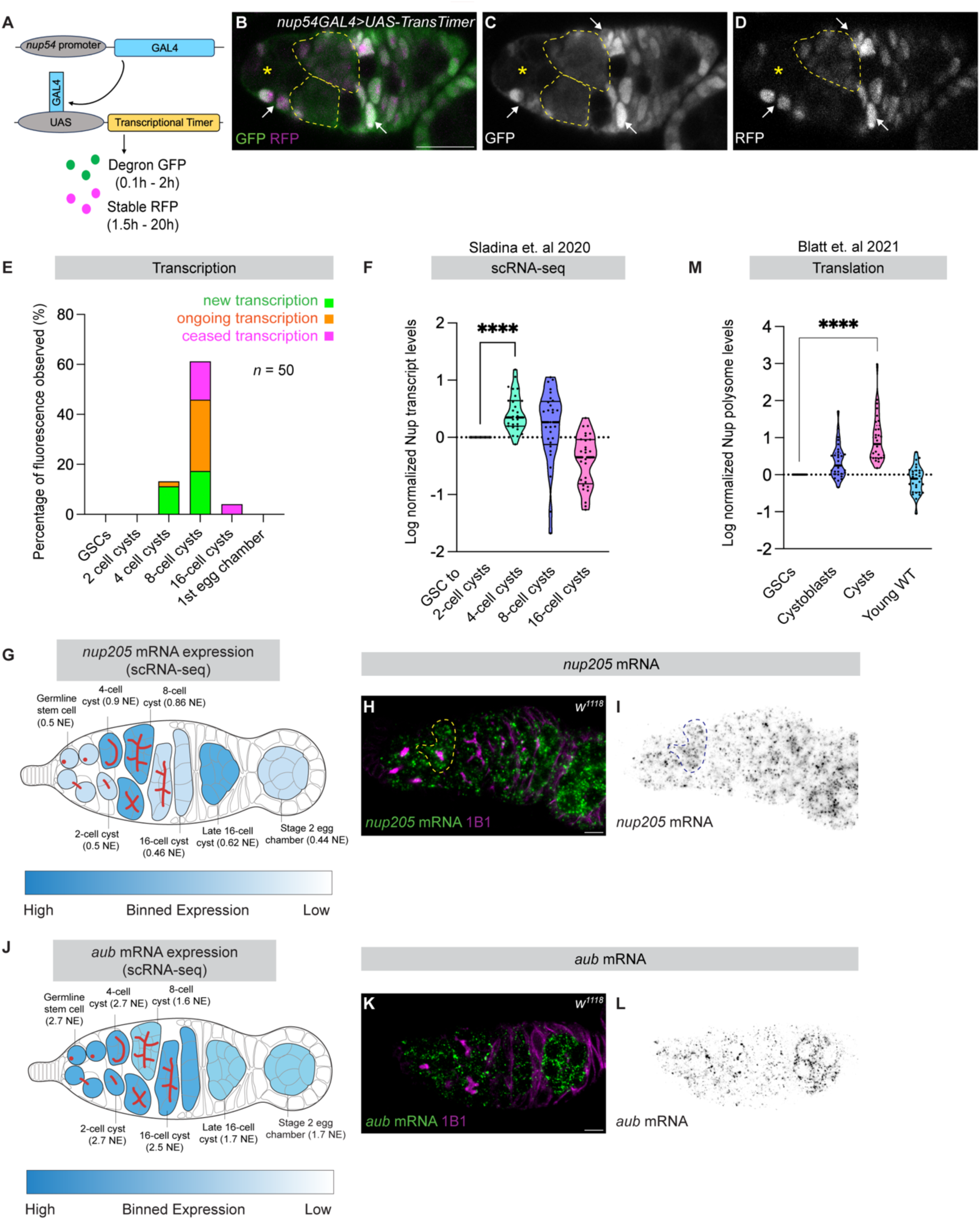
Regulated nucleoporin transcription and translation contribute to dilution of old NPCs. **(A)** Schematic depicting the transcriptional timer. **(B–D)** Confocal images of *nup54GAL4;UAS-TransTimer* germaria showing GFP (new transcription) and endogenous RFP fluorescence (older transcription). Arrows indicate somatic cells exhibiting yellow fluorescence, reflecting continuous *nup54* transcription. Yellow outlines demarcate fluorescence signals in 4-cell cysts (GFP only) and 8-cell cysts (GFP and RFP), respectively. Yellow asterisks mark GSCs, cystoblasts and 2 cell cysts. Scale bar, 25 µm. **(E)** Quantification of GFP and RFP fluorescence patterns from the transcriptional timer reporter across germline stages (*n* = 50 germaria). **(F)** Violin plot showing log-normalized *nup* transcript levels in GSCs and 4-, 8-, and 16-cell cysts from published scRNA-seq datasets. *n* = 29 Nups; **** = *p* < 0.0001. **(G)** Binned expression map of *nup205* mRNA across the germarium derived from scRNA-seq, with darker blue indicating higher transcript abundance. **(H–I)** Confocal images showing in situ detection of *nup205* mRNA in germaria. *nup205* mRNA levels peak during the 4- and 8-cell cyst stages. Scale bar, 5 µm. **(J)** Binned expression map of *aub* mRNA (control) across the germarium from scRNA-seq. **(K–L)** Confocal images showing in situ detection of *aub* mRNA, which remains broadly expressed across germline stages. Scale bar, 5 µm. **(M)** Violin plot showing log-normalized nucleoporin polysome association in GSCs, cystoblasts, differentiating cysts, and young wild-type ovaries, indicating reduced nucleoporin translation during early cyst stages. *n* = 29 Nups; **** = *p* < 0.0001.

To independently validate this timing, we analyzed published single-cell RNA-seq datasets and found that 29 transcripts encoding multiple Nups, including *nup205*, increased beginning at the 4-cell cyst stage (**Fig. 3F–G**) (**Sup. Table 1**) (*4*, *31*, *32*). We confirmed this induction by RNA in situ hybridization for *nup205*, which revealed increased transcript levels during cyst differentiation (**Fig. 3H-I**), in contrast to *aubergine* (*aub*), which exhibited relatively uniform expression across cyst stages (**Fig. 3J-L**). We next asked whether transcriptional induction is accompanied by increased translation by analyzing published polysome-sequencing data (*31*). Translation of 29 Nups was low in undifferentiated cells and increased during cyst stages (**Fig. 3M**) (**Sup. Table 1**), coincident with a known burst of global translation prior to oocyte specification (*33*). Notably, *nup44A*, a Nup required for oocyte fate, is selectively translated during this window (*34*). Together, these findings indicate that Nup transcription and translation are developmentally delayed and activated during cyst differentiation, supporting a model in which dilution of pre-existing NPCs is coupled to the synthesis and incorporation of newly assembled complexes at the onset of oocyte specification.

### ESCRT-III/Vps4 activity is required to reduce NPC levels prior to oocyte specification

Nup transcription and translation peak in 4–8-cell cysts, yet NPC marker signal reaches a minimum at the 8-cell cyst stage. This timing indicates that dilution alone is insufficient to account for the observed reduction in NPC abundance, suggesting that active removal contributes to NPC reduction during this developmental window. In yeast, NPCs are removed from the nuclear envelope by a surveillance pathway involving the ESCRT-III complex and Vps4, which mediates membrane scission and targets NPC components for degradation (**Fig. 4S1A**) (*12*).

**Figure 4:**
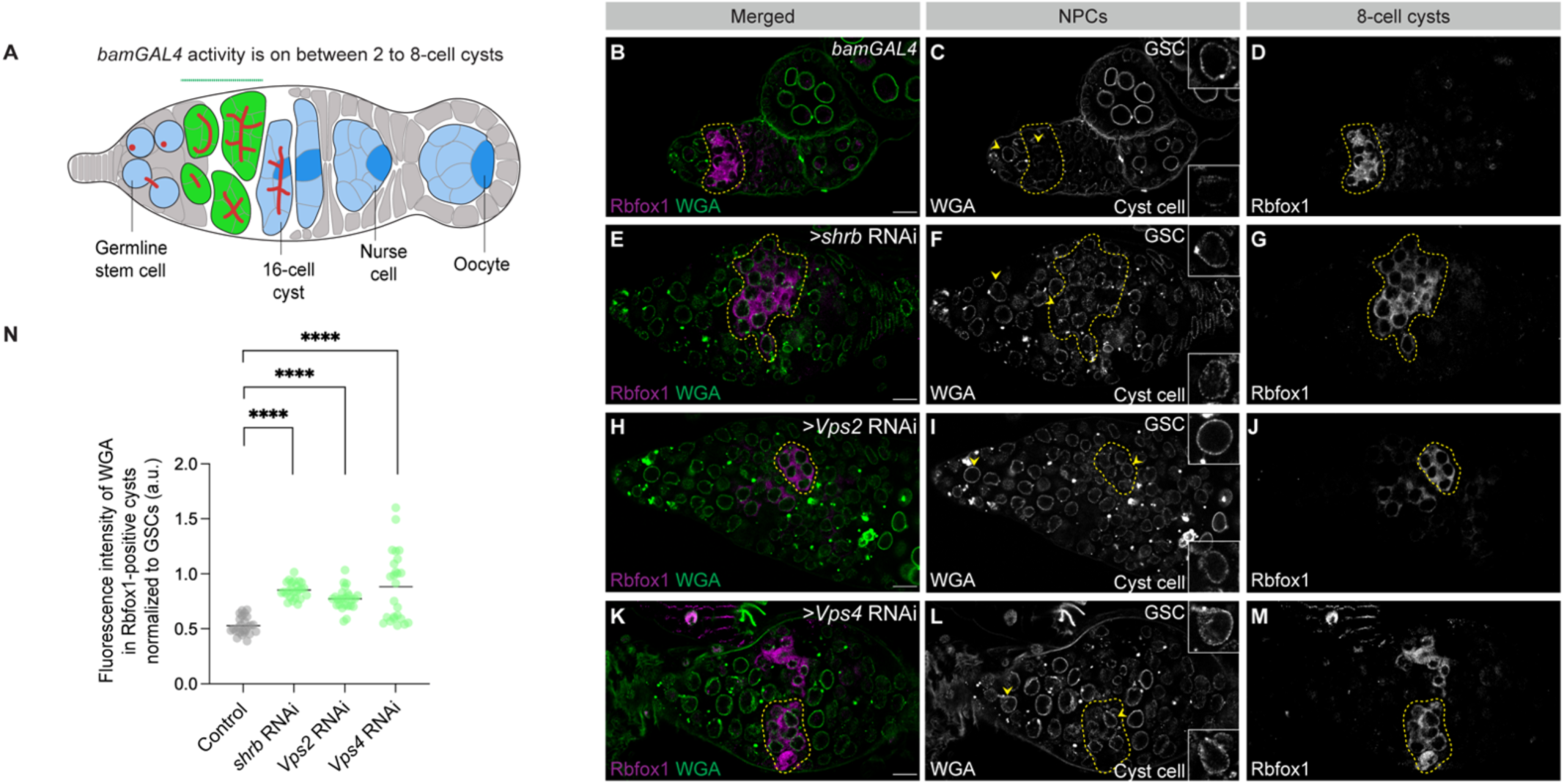
ESCRT-III/Vps4 machinery is required for NPC reduction during cyst differentiation. **(A)** Schematic of the *Drosophila* germarium showing where *bam*GAL4 is active. **(B–M)** Confocal images of control and germline-specific depletion of *shrb*, *Vps2*, or *Vps4* in germaria, stained with Rbfox1 to identify 8-cell cysts and WGA to label NPCs. GSCs are marked with yellow arrowheads, and yellow outlines indicate Rbfox1-positive 8-cell cysts. Scale bars, 10 µm. **(N)** Quantification of WGA fluorescence intensity in Rbfox1-positive cysts, normalized to GSCs within the same germarium, shown in arbitrary units (a.u.). Depletion of *shrb*, *Vps2*, or *Vps4* results in a significant increase in NPC signal relative to control, indicating a failure to reduce NPC levels during cyst differentiation. Statistical analysis was performed using a two-tailed unpaired Welch’s *t*-test. *n* = 5 germaria per genotype; ****, *p* < 0.0001.

To test whether this pathway operates in the Drosophila germline, we depleted key ESCRT-III components, *shrub* (*shrb*), *Vps2*, and *Vps4*, and assessed NPC marker signal during cyst differentiation (*35*). Germline depletion using *nanos*GAL4 resulted in elevated NPC marker signal in Bam-positive cysts, as assessed by mAb414 staining (**Fig. 4S1B–N**). To restrict depletion to differentiating cysts, we used *bam*GAL4 to deplete *shrb*, *Vps2*, or *Vps4* specifically during cyst stages (**Fig. 4A**) (*24*). Quantification of WGA and Elys fluorescence in 8-cell cysts, identified using Rbfox1, revealed a consistent increase in NPC marker signal relative to controls (**Fig. 4B–N**, **Fig. 4S1O–A1**). Importantly, accumulation of 8-cell cysts in *rbfox1* RNAi ovaries did not elevate NPC levels (**Fig. 2G–J**), indicating that increased NPC abundance in ESCRT-depleted cysts is not a secondary consequence of 8-cell cyst accumulation or developmental delay. Together, these results indicate that ESCRT-III/Vps4 activity is required to reduce NPC abundance during cyst differentiation.

In yeast, NPCs removed from the nuclear envelope are delivered to lysosomes for degradation (*36*, *37*). Consistent with engagement of an endolysosomal pathway, we found that Lamp1, a marker of acidic endolysosomal compartments, is expressed in the germline and preferentially translated during cyst stages (**Fig. 4S2A–B**) (*38*). Visualization of acidic compartments using *nanos*GAL4 driven Lamp1::GFP revealed frequent spatial association with NPC markers during the 8-cell cyst stage (**Fig. 4S2C–L**). Together, these observations support a model in which ESCRT-III/Vps4-mediated NPC removal engages the endolysosomal pathway during cyst differentiation.

### ESCRT-III/Vps4 dependent NPC turnover enables germ cell gene silencing and oocyte specification

We previously found that NPCs help maintain stage-specific gene expression programs in the germline (*39*, *40*). In GSCs, proper transcriptional regulation depends on chromatin-organizing proteins and NPCs (*39*), whereas during cyst differentiation, newly synthesized NPCs are required to silence a subset of genes expressed in GSCs (*15*, *40*). We therefore asked whether failure to turn over NPCs during cyst stages leads to persistent expression of germ cell genes. To test this, we examined expression of Blanks, a germ cell gene normally silenced by the 4-cell cyst stage (*4*). As expected, Blanks was expressed in cystoblasts of *bam*-depleted ovaries but was absent from 8-cell cysts in *rbfox1*-depleted ovaries (**Fig. 5S1A–J**). In contrast, depletion of ESCRT-III components (*shrb*, *Vps2*, or *Vps4*) during cyst stages using *bam*GAL4 resulted in persistent Blanks expression in Rbfox1-positive 8-cell cysts (**Fig. 5A–M**). These findings indicate that ESCRT-III/Vps4 activity is required to silence germ cell gene expression during cyst differentiation.

**Figure 5:**
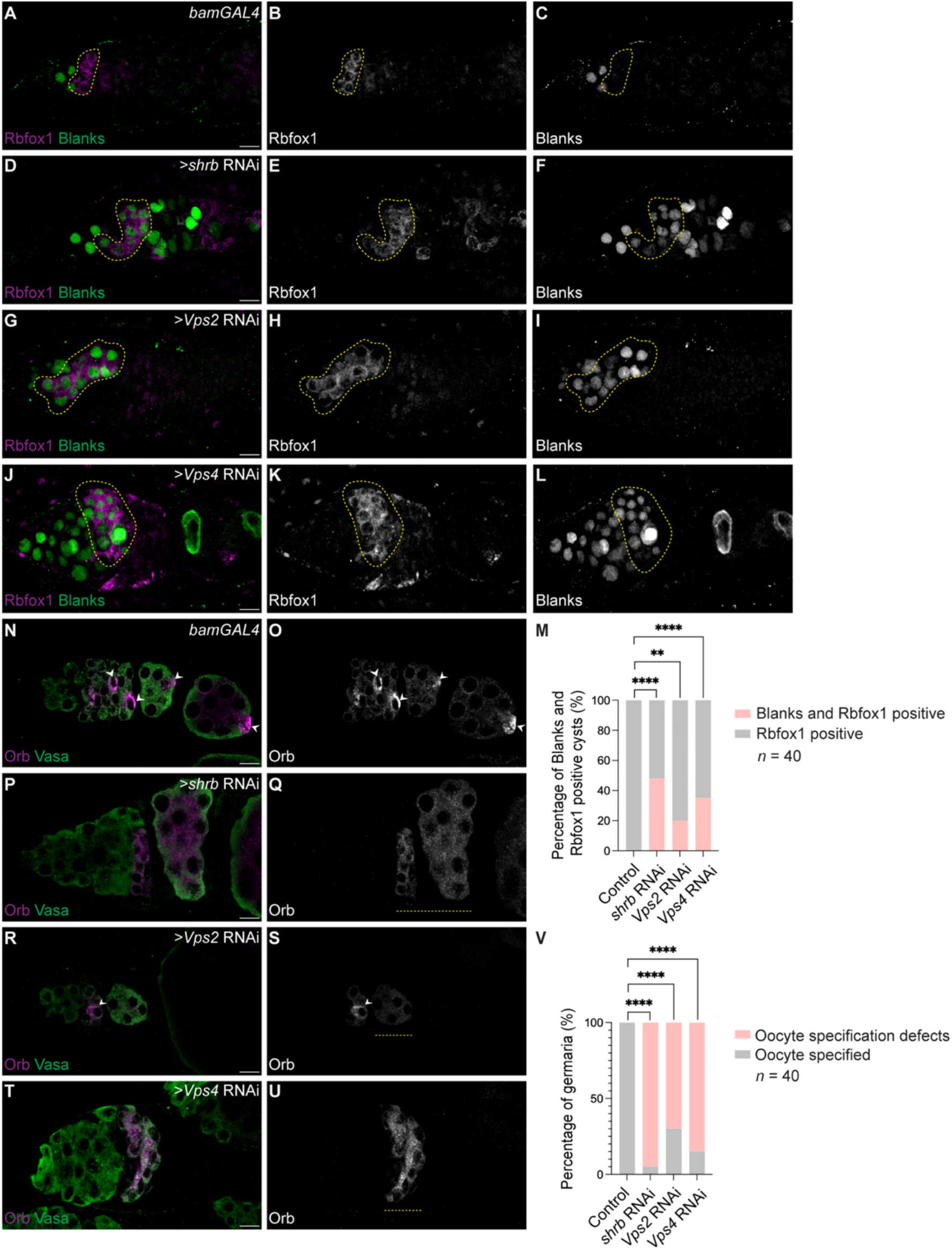
NPC degradation is required for germ cell gene silencing and oocyte specification. **(A–L)** Confocal images of control and germline cyst-specific depletion of *shrb*, *Vps2*, or *Vps4* in germaria, stained for Rbfox1 to identify 8-cell cysts (yellow outlines) and Blanks to mark germ cell gene expression. In control germaria, Blanks expression is extinguished prior to the Rbfox1-positive stage. In contrast, Blanks persists in Rbfox1-positive cysts following depletion of *shrb*, *Vps2*, or *Vps4*. Scale bars, 10 µm. **(M)** Quantification of the percentage of Rbfox1-positive 8-cell cysts that retain Blanks expression in control and *shrb*-, Vps2-, or *Vps4*-depleted ovaries. Statistical analysis was performed using Fisher’s exact test. *n* = 40 germaria per genotype; ****, *p* < 0.0001; **, *p* = 0.0053. **(N–U)** Confocal images of control and *shrb*-, *Vps2*-, or *Vps4*-depleted germaria stained for Orb to assess oocyte specification. In control germaria, Orb localizes to a single cell within the 16-cell cyst (white arrowheads), corresponding to the presumptive oocyte. In contrast, Orb remains distributed across multiple cells in ESCRT-III/Vps4-depleted germaria (yellow dashed outlines), indicating defective oocyte specification. Some cysts show defective oocyte specification in upon depletion of *Vps2*. Scale bars, 10 µm. **(V)** Quantification of the percentage of germaria exhibiting a specified oocyte based on Orb localization in control and *shrb*-, *Vps2*-, or *Vps4*-depleted ovaries. Statistical analysis was performed using Fisher’s exact test. *n* = 40 germaria per genotype; ****, *p* < 0.0001.

Because germ cell gene silencing is tightly linked to oocyte fate (*15*), we next tested whether ESCRT-III/Vps4-dependent NPC turnover is required for meiotic entry and oocyte specification. In wild-type ovaries, synaptonemal complex assembly initiates in 8-cell cysts, as marked by crossover suppressor on 3 of Gowen (c(3)G) staining, and becomes restricted to a single cell by the 16-cell cyst stage, corresponding to the future oocyte (**Fig. 5S1K–L**) (*41*). In contrast, ESCRT-III/Vps4-depleted ovaries largely lacked c(3)G-positive cells in 8-cell cysts, indicating a defect in synaptonemal complex assembly (**Fig. 5S1M–S**). Consistent with disrupted oocyte specification, Orb was expressed but failed to localize to a single cell in 16-cell cysts in ESCRT-III/Vps4-depleted ovaries (**Fig. 5N–V**). Together, these findings strongly suggests that ESCRT-III/Vps4-dependent NPC turnover is required to extinguish germ cell gene expression, enable meiotic recombination, and establish oocyte identity during cyst differentiation.

### Cyst-specific NPC reduction rescues gene expression defects caused by ESCRT-III depletion

To test whether compromised germ cell gene silencing following ESCRT-III depletion results from persistent NPCs, we asked whether targeted reduction of NPCs could rescue this phenotype. To reduce NPC abundance specifically during cyst stages, we depleted *torsin*, which is known to destabilize NPCs by disrupting their assembly and maintenance (*42*, *43*). For spatial control, *bam*GAL4 was used to drive both *shrb* RNAi and *torsin* RNAi in differentiating cysts.

In wild-type cysts, Blanks expression is extinguished during differentiation (**Fig. 6A–C, M**). Depletion of the ESCRT-III component *shrub* elevated NPC marker signal and caused persistent Blanks expression (**Fig. 6D–F, M, 6S1A-F, 6S1M**), reflecting failure to silence germ cell gene expression. *torsin* RNAi also had reduced NPC marker signal in the cyst stages (**Fig. 6S1G-I, 6S1M**) and did not induce persistent Blanks expression (**Fig. 6G–I, M**), indicating that this degree of NPC reduction is not sufficient to disrupt germ cell gene silencing. Co-depletion of *torsin* and *shrub* lowered NPC marker signal relative to *shrub* RNAi alone (**Fig. 6S1J–L, M**) and suppressed persistent Blanks expression (**Fig. 6J–M**). Consistent with restored germ cell gene silencing, meiotic recombination and Orb restriction to a single cell was partially rescued (**Fig. 6N-V, 6S1Q-Z**). Co-expression of an unrelated UAS-GFP transgene failed to rescue oocyte specification defects in *shrub* RNAi (**Fig. 6S2A–M**), demonstrating that suppression is specific to NPC reduction rather than increased GAL4/UAS transgene load.

**Figure 6:**
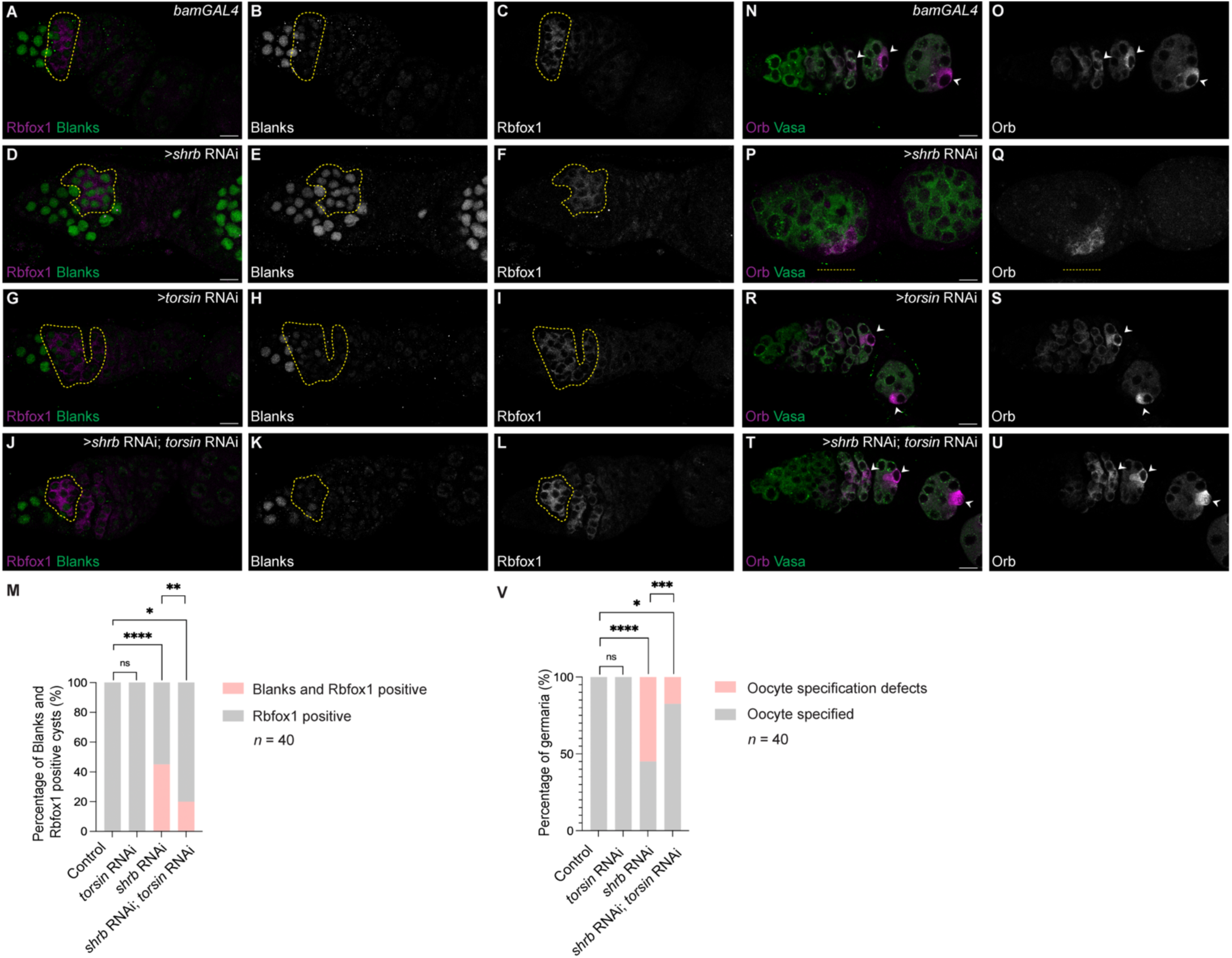
Forced NPC reduction during cyst differentiation rescues gene expression and oocyte specification defects caused by loss of ESCRT III component, Shrb. **(A–I)** Confocal images of control, *shrb*-, and *torsin*-depleted germaria stained for Rbfox1 and Blanks to assess termination of germ cell gene expression. **(J–L)** Confocal images of *shrb* and *torsin* double-depleted germaria showing timely extinction of Blanks expression. **(M)** Quantification of the percentage of Rbfox1-positive 8-cell cysts retaining Blanks expression in control, *shrb*-, *torsin*-, and *shrb*/*torsin* double-depleted ovaries. Statistical analysis was performed using Fisher’s exact test. *n* = 40 germaria per genotype; ns, *p* > 0.9999; *, *p* = 0.0307; **, *p* = 0.0053; ****, *p* < 0.0001. **(N–S)** Confocal images of control, *shrb*-, and *torsin*-depleted germaria stained for Orb and Vasa to assess oocyte specification. White arrowheads indicate the specified oocyte. **(T–U)** Confocal images of germaria with combined depletion of *shrb* and *torsin*. Oocyte specification is restored upon forced NPC reduction during cyst stages induced by *torsin* depletion. **(V)** Quantification of the percentage of germaria exhibiting a specified oocyte in control, Shrb-, Torsin-, and Shrb/Torsin double-depleted ovaries. Statistical analysis was performed using Fisher’s exact test. *n* = 40 germaria per genotype; ns, *p*> 0.9999; *, *p* = 0.0117; ***, *p* = 0.0010; ****, *p* < 0.0001. Yellow dashed outlines mark Rbfox1-positive 8-cell cysts. Scale bars, 10 µm.

### A subset of NPCs localizes to acidic endolysosomal compartments in mouse oocytes

To assess whether NPC turnover may also occur in mammals, we examined NPC localization in sexually mature mouse oocytes, which enter meiosis during fetal development and remain arrested in prophase I for extended periods (*44*, *45*). This prolonged arrest raises the possibility that long-lived nuclear structures such as NPCs may undergo turnover during oocyte maturation. Immunostaining of prophase I–arrested oocytes with the NPC marker mAb414 revealed robust signal at the nuclear envelope, consistent with intact NPCs, as well as discrete mAb414-positive puncta in the cytoplasm (**Fig. 6S3A–C**). A subset of these puncta was unusually large (∼4 μm in diameter; n = 6 oocytes) and morphologically resembled late endolysosomal compartments.

To test whether these NPC-containing structures were associated with acidic endolysosomal compartments, we co-stained oocytes for mAb414 and Lamp1 (*46*, *47*). Nearly all large Lamp1-positive aggregates overlapped with cytoplasmic mAb414 signal (**Fig. 6S3D–F**), indicating that a subset of NPC components localize to Lamp1-positive compartments in mouse oocytes. These observations are consistent with endolysosomal-associated NPC turnover in mammalian oocytes, analogous to the pathway identified in Drosophila.

## Discussion

The germline must renew itself each generation, preserving nuclear function while discarding components assembled in earlier developmental contexts. Here, we identify NPC replacement as a developmentally programmed mechanism required for oocyte specification in Drosophila. During cyst differentiation, NPCs assembled during earlier germ cell stages are actively removed by the ESCRT-III/Vps4 pathway, while Nup transcription and translation are reactivated to drive de novo NPC assembly. Disrupting this coordinated cycle of NPC removal and replacement results in persistent germ cell gene expression, defective meiotic entry, and failure of oocyte specification, establishing NPC turnover as a functionally essential nuclear remodeling event during the germ cell-to-maternal transition (**Fig. 6S3G**).

Our findings further suggest that NPC turnover represents a conserved strategy for nuclear renewal in the germline. In *Saccharomyces cerevisiae*, NPC subunits assembled prior to sporulation are selectively eliminated to ensure inheritance of newly synthesized NPCs (*48*). Consistent with this principle, we observe NPC components associated with acidic endolysosomal compartments in mouse oocytes during prolonged prophase I arrest, a stage in which oocytes remain metabolically active for months to years. Although the molecular machinery underlying NPC clearance in mammals remains to be defined, these observations raise the possibility that endolysosomal-associated NPC turnover is a conserved feature of gametogenesis.

NPCs are among the most stable macromolecular assemblies in the cell, yet their longevity renders them vulnerable to cumulative damage, as evidenced by NPC deterioration in aging cells and neurodegenerative disease (*11*). We propose that NPC turnover during oogenesis functions as a nuclear rejuvenation mechanism, limiting the inheritance of potentially compromised NPCs and ensuring proper chromatin organization and transcriptional control in the embryo. More broadly, our findings also support a model in which nuclear envelope remodeling contributes directly to cell fate transitions by resetting nuclear architecture. Given the parallels between oocytes and other long-lived or post-mitotic cells, NPC clearance pathways may represent a general strategy to preserve nuclear integrity during development, aging, and stress.

## Materials and Methods

### Fly lines

The following RNAi and mutant fly stocks were used in this study: *bam* RNAi (Bloomington Drosophila Stock Center [BDSC], 58178), *rbfox1* RNAi/CyO (the Buszczak Laboratory), *shrb* RNAi (BDSC, 38305); *Vps2* RNAi (Vienna Drosophila Resource Center [VDRC] v24869); *Vps4* RNAi (VDRC, v35126).

The following tagged line was used in this study: *UAS-dcr2i;nosGAL4;bam::GFP* (Lehmann Laboratory); *nup107::GFP*(BDSC 35514), *nup358::GFP* (Beck Laboratory), Fs(2)*Ket-GFP/CyO* (BDSC 35515); *nup358::E5* timer (this study, WellGenetics), and *tnpo-SR::mCherry* (Ables Laboratory) (*27*).

The germline-specific drivers and double-balancer lines used in this study were *UAS-dcr2i;nosGAL4* (BDSC 25751), *bamGAL4* (BDSC 80579), *nup54GAL4* (BDSC 80664); *nosGAL4;MKRS/TM6* (BDSC 4442) and *If/CyO;nosGAL4*.

### Generation of the Nup358-E5 timer fly line

The Nup358-E5 timer fly line was generated by WellGenetics via CRISPR/Cas9-mediated genome editing by homology-dependent repair (HDR) using a guide RNA and a dsDNA plasmid donor. Cassette DsRed-E5-PBacw was inserted right after +15 nt from ATG of *nup358*. The cassette contains Fluorescence Timer DsRed-E5 (*29*) and a selection marker PBacw. DsRed-E5 is a tetrameric fluorescent timer protein that changes its fluorescence from over time. It is an artificial derivative of the naturally occurring fluorescent protein encoded by the Discosoma drFP583 gene (GenBank:AF168419; AAF03369), derived by mutagenesis of DsRed1. It contains the mutations V105A and S197T relative to DsRed1. PBacw contains Piggy Bac 3’ terminal repeats, hsp70 promoter, *white*CDS and Piggy Bac 5’ terminal repeats. After the cassette was injected into embryos from the w[1118] fly strain, genetic screening and crosses were performed with the progeny. The selection marker was then excised by Piggy Bac transposase. Only one TTAA motif was left from the cassette after transposition. PAM mutations were incorporated into the edited genome to make the donor inactive to guide RNA. Finally, the genomic sequence of the nup358-E5 timer fly was confirmed by performing PCR and Sanger sequencing.

### Reagents for fly husbandry

Fly crosses were grown at 25°C–29°C and dissected between 0 and 3 days after eclosion. Fly food for stocks and crosses was prepared using the published laboratory protocol (summer/winter mix), and narrow vials (Fisherbrand Drosophila vials, Fisher Scientific) were filled to ∼10–12 mL.

### Dissection and immunostaining

Ovaries were dissected, and the ovarioles were separated using mounting needles in PBS solution and kept on ice. Samples were then fixed for 10 min in 10% methanol-free formaldehyde. Ovaries were washed in 0.5 mL of PBT (1× PBS, 0.5% Triton X-100, 0.3% BSA) four times for 10 min each while incubating on a nutator. Primary antibodies in PBT were added and incubated overnight at 4°C while mixed on the nutator. Samples were next washed three times for 10 min each in 0.5 mL of PBT. One last wash was performed for 10 min in 0.5 mL of PBT with 2% donkey serum. Secondary antibodies were added in PBT with 4% donkey serum and incubated for 3 - 4 h at room temperature. When required, WGA was also added at this step at a dilution of 1:500 along with secondary antibodies in PBT with 4% donkey serum and incubated for 3 - 4 h at room temperature. Samples were washed three times for 10 min each in 1 mL of 1× PBST (0.2% Tween 20 in 1× PBS) and incubated in VectaShield or VectaShield Plus with DAPI (Vector Laboratories) for at least 2 h before mounting.

For NPC staining, the above protocol was performed with the following changes: The eppendorf tube was coated with 3% BSA before fixing. Samples were fixed for 10-15 min in 600 uL of 5% PFA in PBS. Ovaries were washed with 0.5ml of PBT (1× PBS, 0.3% Triton X-100) four times for 10 min each while incubating on a nutator. Primary antibodies in PBT were added and incubated overnight at 4°C while mixing on a nutator. Chicken anti-GFP (from Aves Labs) in this context was used with a dilution of 1:250. Samples were next washed three times for 5 min each in 1 mL of PBT. Secondary antibodies were added in PBT with a dilution of 1:750 in this context and incubated for 2 h at room temperature. Samples were washed four times for 5 min each in PBST (1× PBST, 0.2% Tween-20). Ovaries were incubated in VectaShield (or VectaShield Plus with DAPI (Vector Laboratories) for at least 2 h. Ovaries were then transferred onto mounting slides.

The primary antibodies used were rabbit anti-Elys (1:1000; Capelson laboratory), anti-Nup153 (1:500; Capelson laboratory), rabbit anti-Nup98 (1:1000; Capelson laboratory), guinea pig anti-Rbfox1 (the Buszczak Laboratory), mouse anti-1B1 (1:20; Developmental Studies Hybridoma Bank [DSHB]), rabbit anti-Vasa (1:1000; Rangan laboratory, Flora et al., 2018), chicken anti-Vasa (1:1000; Rangan laboratory, Flora et al., 2018), rabbit anti-GFP (1:2000; Abcam ab6556), chicken anti-GFP (1:2000; Abcam ab13970), mouse anti-Orb (1:30; DSHB; 4H8), mouse anti-NPC (1:150; BioLegend AB_2565026), rabbit anti-blanks (1:1000, the vSontheimer laboratory), rabbit anti-PH3 (1:200, Cell Signaling 9701), mouse anti-lamin B (1:5, DSHB, ADL195), mouse anti-C(3)G (1:500, the Hawley laboratory). The following secondary antibodies were used: Alexa 488 (1:500; Molecular Probes), Cy3 (1:500; Jackson Laboratories), Cy5 (1:500; Jackson Laboratories), anti-WGA-Alexa 488 (1:500; ThermoFisher Scientific), and anti-WGA-Alexa 594 (1:500; ThermoFisher Scientific).

### Fluorescence imaging

Ovaries were mounted on slides and imaged using Zeiss LSM-880 and LSM-980 confocal microscopes under 20×, 40×, and 63× oil objectives with pinhole set to one airy unit. Image processing was done using Fiji, and gain adjustment and cropping were performed in Adobe Photoshop 25.6. All colocalization analysis and quantification for determining NPC levels in oocytes and nurse cells were performed using Imaris 10.2.

### RNA in situ hybridization

All steps were done using RNAse free reagents and supplies with gentle rotation, except for steps at 37°C. The protocol was adapted from Choi et al (*49*). Specimens were fixed in PBS, 0.1% Tween-20 (Tw), and 4% methanol-free formaldehyde for 20 min at room temperature; washed twice with PBS and 0.1% Tw at room temperature; and dehydrated with sequential washes with 25%, 50%, 75%, and 100% ethanol in PBS for 5 min each on ice. Samples were stored at least overnight (up to 1 wk) in 100% ethanol at −20°C. Samples were rehydrated with sequential washes with 100%, 75%, 50%, and 25% ethanol in PBS on ice; permeated for 2 h in PBS and 1% Triton X-100 at room temperature; postfixed in PBS, 0.1% Tw, and 4% paraformaldehyde for 20 min at room temperature; washed twice with PBS and 0.1% Tw for 5 min on ice; washed with 50% PBS and 0.1% Tw/50% 5× SSCT (5× SSC, 0.1% Tween) for 5 min on ice; washed twice with 5× SSCT for 5 min on ice; incubated in probe hybridization buffer for 5 min on ice; prehybridized in probe hybridization buffer for 30 min at 37°C; and hybridized overnight (16–24 h) at 37°C. Probe concentrations were determined empirically, and ranged from 8 to 16 pmol of each probe in 1 mL; probe solution was prepared by adding probes to prewarmed probe hybridization solution. After hybridization, specimens were washed four times with probe wash buffer for 15 min each at 37°C, and twice with 5× SSCT for 5 min each at room temperature. Specimens were equilibrated in amplification buffer for 5 min at room temperature. Hairpin solutions were prepared by heating 30 pmol of each hairpin for 90 s at 95°C, cooling at room temperature in the dark for 30 min and subsequently adding the snap-cooled hairpins to 500 μL of amplification buffer at room temperature. Specimens were incubated in hairpin solution overnight (∼16 h) at room temperature and washed multiple times with 5× SSCT—twice for 5 min, twice for 30 min, and once for 5 min. DAPI was added in the first 30 min wash. Specimens were equilibrated in VectaShield overnight at 4°C and mounted in VectaShield or further stained using the immunofluorescence protocol (see above).

### Mouse oocyte collection, immunostaining, and imaging

Prophase I-arrested oocytes were isolated from ovaries of 6-week-old CF1 females (Envigo) injected 48 h earlier with 5 I.U. of pregnant mare’s serum gonadotropin (PMSG) (Lee Biosolutions #493–10). Oocytes were collected as described previously. Briefly, ovaries were placed in minimal essential medium (MEM) containing 2.5 μM milrinone (Sigma-Aldrich #M4659) to prevent meiotic resumption and oocytes from antral follicles were isolated by piercing the ovaries with needles. After treating oocytes with Acidic Tyrode’s solution (Millipore Sigma #MR-004-D) to remove the *zona pellucida*, oocytes were fixed in 4% PFA in PBS for 1h at room temperature, followed by permeabilization in PBS containing 0.2% Triton X-100 for 20 min and blocking in 0.3% BSA containing 0.01% Tween in PBS for 10 min. Immunostaining was performed by incubating in primary antibodies: mAb414 (1:1000; mouse) to detect the nuclear pore, LAMP1 (1:100; rabbit; Abcam #ab24170) to detect lysosomes. Antibody incubation occurred for 1-2 h in a dark, humidified chamber, followed by three washes of 10 min each in blocking solution. Oocytes were then incubated in secondary antibodies: Alexa Anti-mouse 568 (1:200; Invitrogen, A10037) and Alexa Anti-rabbit 647 (1:200; Invitrogen, A34573) for 1h in a dark humidified chamber, followed by three washes of 10 min each in blocking solution. After washing, oocytes were mounted in 5 μl of Vectashield containing 4, 6-Diamidino-2-Phenylindole, Dihydrochloride (DAPI) (Life Technologies #D1306). Images were captured using a Leica SP8 confocal microscope equipped with a 40X 1.30 NA oil immersion objective. Optical Z-stacks of 1 μm step with a zoom of 2 were taken.

Mice were housed on a 12/12-h light–dark cycle, with constant temperature and with food and water provided ad libitum. Animals were maintained in accordance with guidelines of the Institutional Animal Use and Care Committee of Rutgers University (protocol 201702497

## Acknowledgements

We thank members of the Rangan Laboratory for insightful discussions and feedback on this project. We are grateful to Drs. Daria Siekhaus and Pooja Flora for critical reading of the manuscript, and to Drs. Michael Buszczak, Elizabeth Ables, Scott Hawley, Erik J. Sontheimer, Maya Capelson, and Martin Beck for reagents and resources.; Nikolaos Tzavaras, Glenn Doherty and Arun Narasimhan from Mount Sinai’s Microscopy and Advanced Bioimaging CoRE for their assistance with confocal imaging and Imaris analysis; Bloomington Drosophila Stock Center, Vienna Drosophila Resource Center, FlyBase for transgenic fly stocks.

## Funding

This work was supported by the National Institutes of Health grants R35 GM156185 to PR; R35 GM136340-06 to KS; R01 GM124143 to MC, R56 AG082906 to PR and MC.

## Author contributions

Conceptualization: PR, MC, KS, SV. Methodology: PR, SV, MC, KS. Investigation and Visualization in *Drosophila*: SV, TN, MP. Investigation and Visualization in the mouse: CB. Funding acquisition: PR, MC, KS. Supervision: PR, MC. Writing - original draft: PR, SV. Writing - reviewing and editing: PR, SV, MC.

## Competing interests

The authors declare no competing interests.

## Data and materials availability

No new code was generated for this work; all analyses have been performed according to the details described in the Methods section. Reagents generated in this study are available from the corresponding author upon reasonable request and completion of a material transfer agreement.

---

**Figure 1S1.**
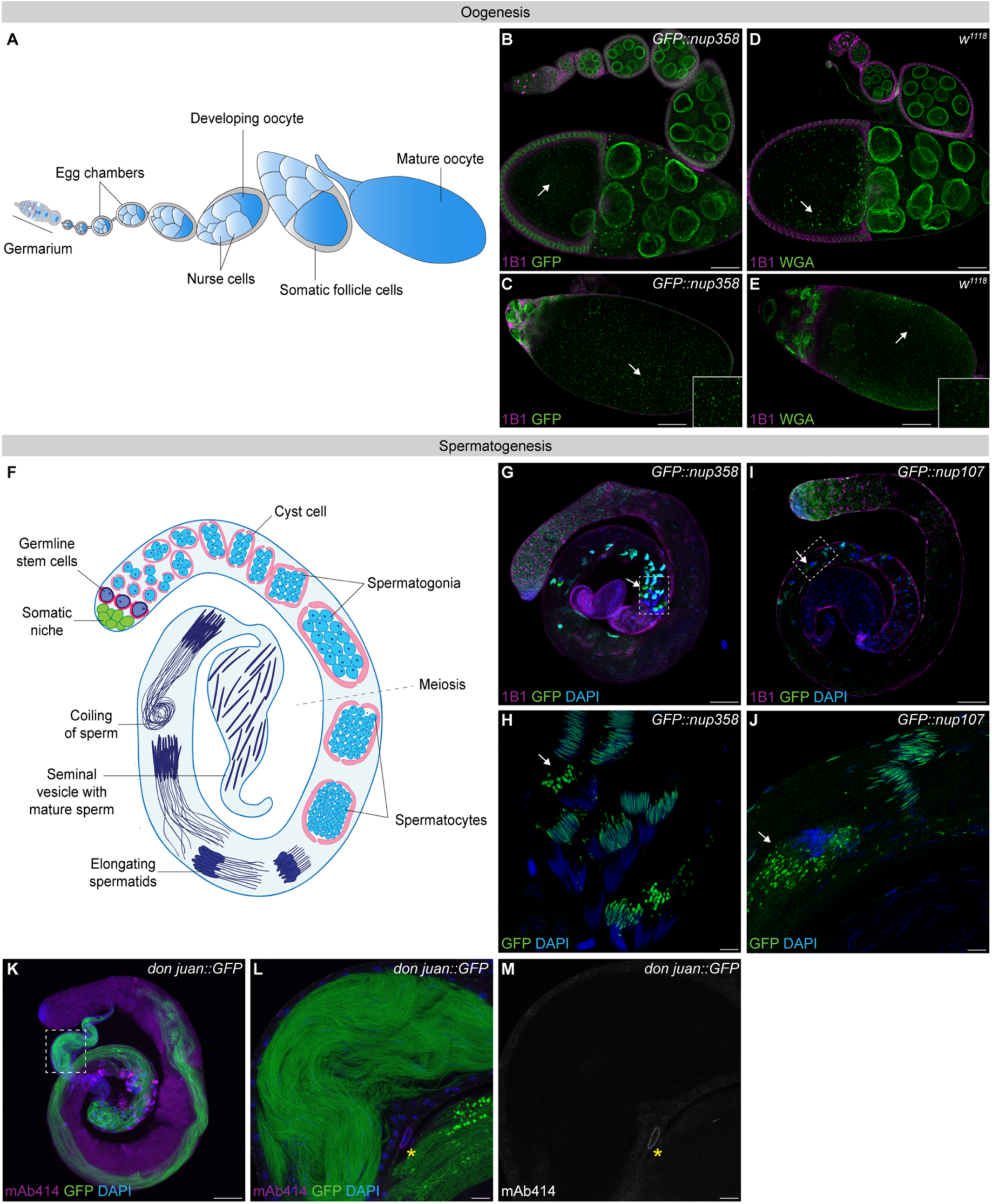
NPC and nucleoporin marker signal accumulates in oocytes but is largely absent from mature sperm. **(A)** Schematic of the *Drosophila* ovariole. **(B)** Confocal image of a *GFP::nup358* ovariole stained for 1B1 and GFP. Arrow indicates accumulation of GFP::Nup358 signal within the ooplasm of the developing oocyte. **(C)** Confocal image of a stage 12 egg chamber showing accumulation of GFP::Nup358 in the ooplasm (arrow). **(D)** Confocal image of a *w*¹¹¹⁸ ovariole stained for 1B1 and wheat germ agglutinin (WGA); arrow indicates accumulation of NPC marker signal in the ooplasm of the developing oocyte. **(E)** Confocal image of a stage 11 egg chamber showing ooplasmic accumulation of NPC marker signal (arrow). **(F)** Schematic of the *Drosophila* testis. **(G)** Confocal images of testis expressing *GFP::nup358*. White dotted boxes indicate late-stage spermatids in which the nucleoporin signal is not associated with DNA. **(H)** Enlarged inset from (G) showing loss of GFP::Nup358 signal from late spermatids (arrow). Scale bar, 10 µm. **(I)** Confocal images of testis expressing *GFP::nup107*. White dotted boxes indicate late-stage spermatids in which the nucleoporin signal is not associated with DNA. **(J)** Enlarged inset from (I) showing loss of GFP::Nup107 signal from late spermatids (arrow). Scale bar, 10 µm. **(K)** Confocal image of *don juan::GFP* testis stained for GFP and mAb414. White dotted box indicates the seminal vesicle containing mature sperm. **(L)** Enlarged inset from (K) showing mature sperm in the seminal vesicle lacking detectable NPC marker signal. Scale bar, 10 µm. **(M)** Grayscale channel illustrating absence of mAb414 signal in mature sperm; yellow asterisk marks a neighboring somatic cell serving as an internal positive control for NPC marker staining. Scale bars, 100 µm unless otherwise indicated.

**Figure 1S2.**
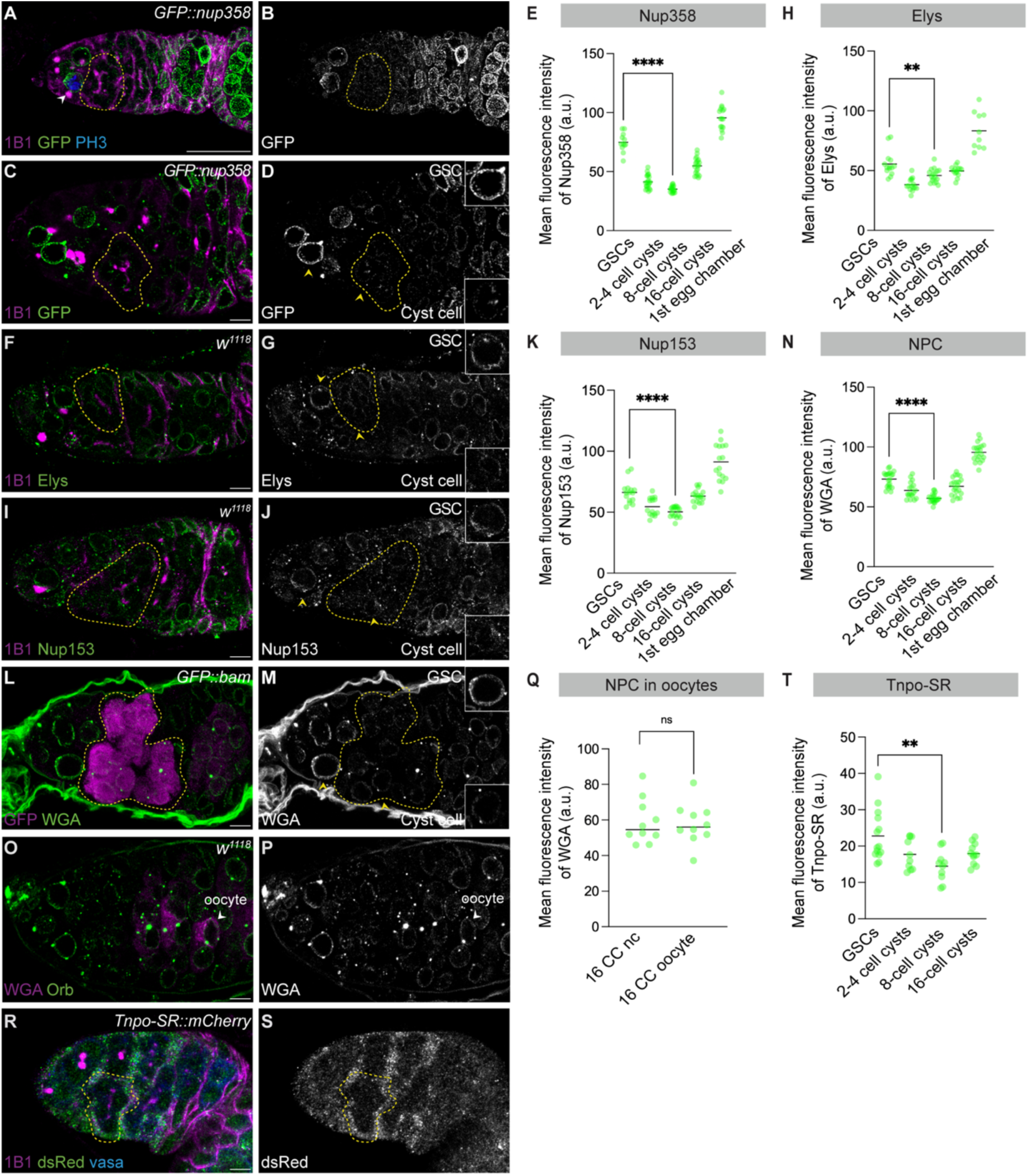
NPC marker signal is reduced during cyst differentiation independent of cell division and nuclear envelope breakdown. **(A–B)** Confocal images of *GFP::nup35*8 germaria. Scale bar, 25 µm. **(C–D)** Confocal images of *GFP::nup358* germaria highlighting NPC marker signal in GSCs and cyst cells. Scale bar, 5 µm. **(E)** Quantification of mean GFP::Nup358 fluorescence intensity (a.u.). **(F–G)** Confocal images of *w*¹¹¹⁸ germaria stained for Elys. Scale bar, 5 µm. **(H)** Quantification of mean Elys fluorescence intensity (a.u.); *n* = 5 germaria; *n* = 5 germaria; ** = *p* = 0.0057. **(I–J)** Confocal images of *w*¹¹¹⁸ germaria stained for Nup153. Scale bar, 5 µm. **(K)** Quantification of mean Nup153 fluorescence intensity (a.u.). **(L–M)** Confocal images of *bam::GFP* germaria showing NPC marker signal in differentiating cysts. Scale bar, 5 µm. **(N)** Quantification of mean WGA fluorescence intensity (a.u.). **(O–P)** Confocal images of *w*¹¹¹⁸ germaria stained for the oocyte marker Orb; arrowheads indicate oocytes. Scale bar, 5 µm. **(Q)** Quantification of mean WGA fluorescence intensity in oocytes (a.u.); *n* = 5 germaria; ns = *p* = 0.8258. **(R–S)** Confocal images of *Tnpo-SR::mCherry* germaria. Scale bar, 5 µm. **(T)** Quantification of mean Tnpo-SR fluorescence intensity (a.u.); *n* = 5 germaria; ** = *p* = 0.0015. Yellow dotted lines indicate 8-cell cysts. Statistical analysis where applicable was performed using a two-tailed unpaired Welch’s *t-*test on mean fluorescence intensity. *n* = 5 germaria; **** = *p* < 0.0001 unless otherwise indicated.

**Figure 3S1:**
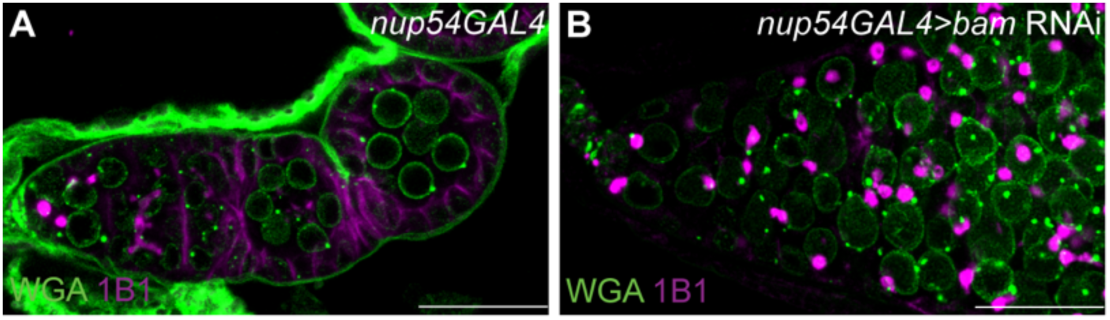
Nup transcription is active in the GSCs. **(A-B)**Confocal images of *nup54GAL4* (control) germaria and *nup54* GAL4 driving *bam* RNAi. RNAi germaria phenocopy germline *bam* depletion, showing accumulation on GSCs/cystoblasts. This indicates that the *nup54* promoter is active in the GSCs/cystoblasts. Scale bars: 25 µm.

**Figure 4S1:**
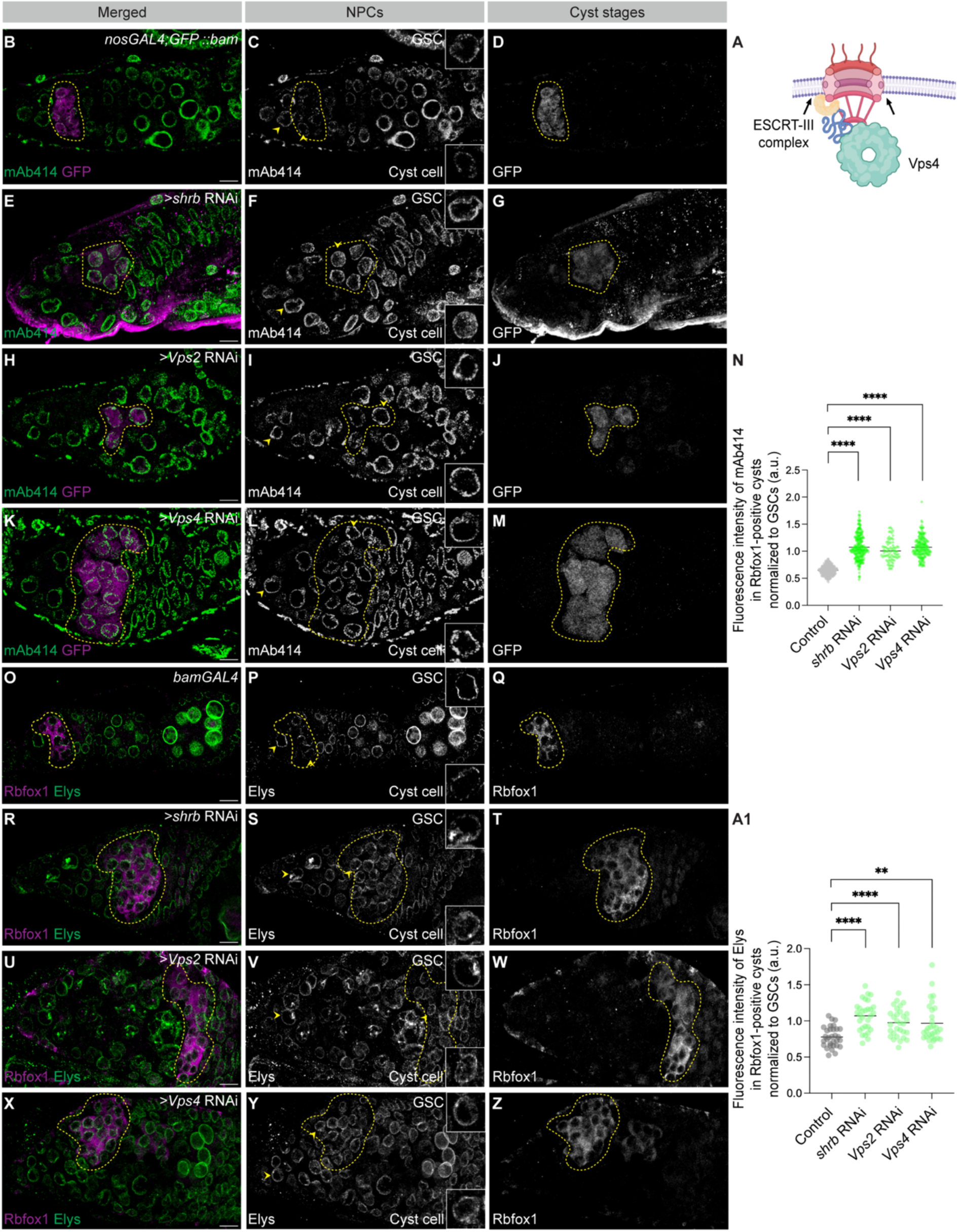
ESCRT-III/Vps4 machinery mediates reduction of nucleoporin levels during cyst differentiation. **(A)** Schematic representation of the proposed ESCRT-III/Vps4-dependent pathway for NPC degradation. **(B–M)** Confocal images of control (*nosGAL4;GFP::bam*) and germline-specific depletion of *shrb*, *Vps2*, or *Vps4* in germaria, stained with mAb414 to label FG-repeat nucleoporins. GFP marks differentiating cysts. GSCs are indicated by yellow arrowheads, and yellow outlines denote 8-cell cysts. Scale bars, 10 µm. **(N)** Quantification of mAb414 fluorescence intensity in GFP-positive cysts, normalized to GSCs within the same germarium and expressed in arbitrary units (a.u.). Depletion of *shrb*, *Vps2*, or *Vps4* results in a significant increase in nucleoporin signal relative to control. Statistical analysis was performed using a two-tailed unpaired Welch’s *t*-test. *n* > 5 germaria per genotype; ****, *p* < 0.0001. **(O–Z)** Confocal images of control and *shrb*-, *Vps2*-, or *Vps4*-depleted germaria stained for Elys. GSCs are indicated by yellow arrowheads, and yellow outlines mark Rbfox1-positive 8-cell cysts. Scale bars, 10 µm. **(A1)** Quantification of Elys fluorescence intensity in Rbfox1-positive cysts, normalized to GSCs within the same germarium and expressed in arbitrary units (a.u.). Depletion of *shrb*, *Vps2*, or *Vps4* leads to a significant increase in Elys signal compared to control. Statistical analysis was performed using a two-tailed unpaired Welch’s *t*-test. *n* = 5 germaria per genotype; ****, *p* < 0.0001; **, *p* = 0.0012.

**Fig. 4S2:**
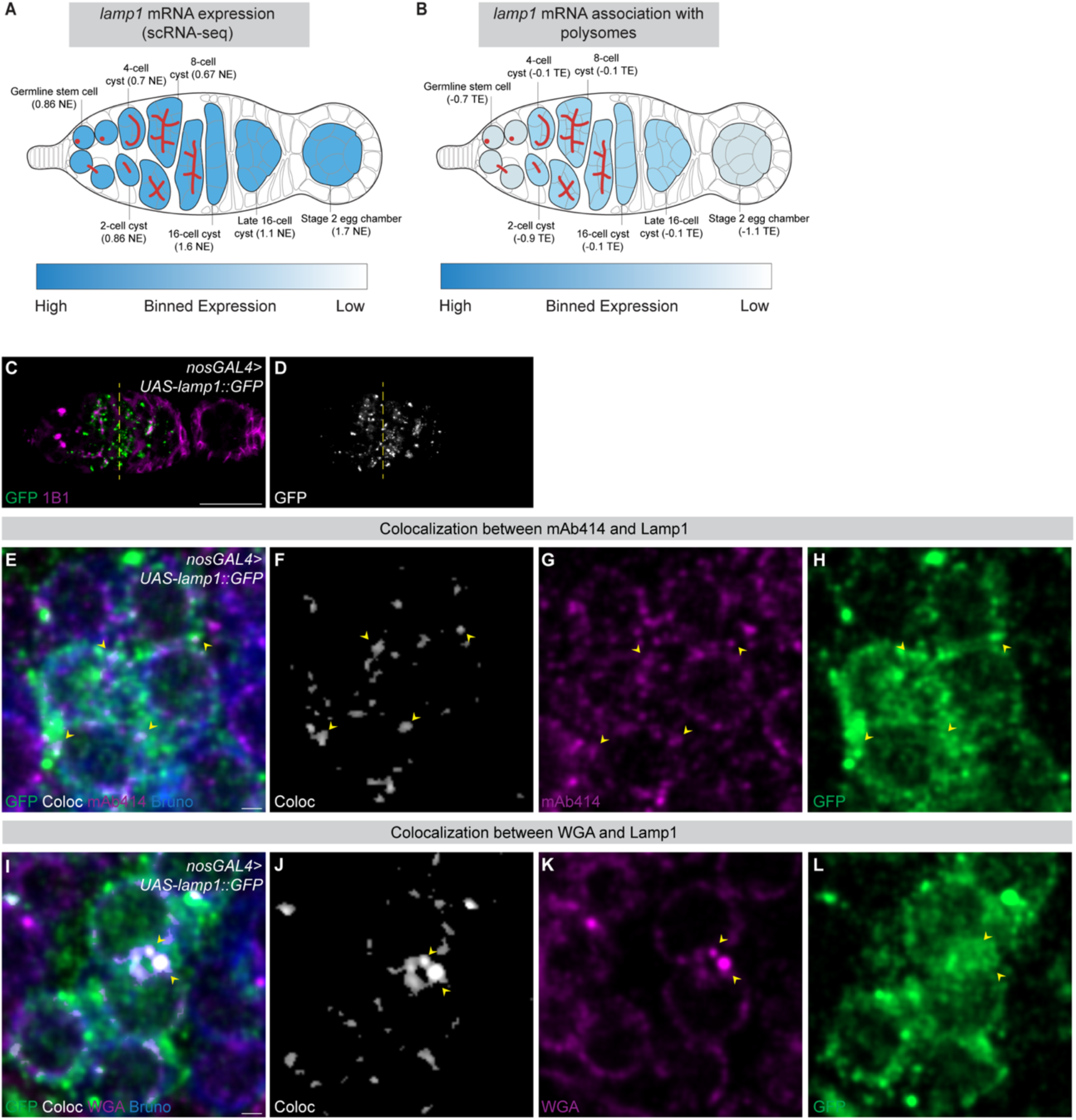
Nucleoporins associate with Lamp1-positive vesicles during cyst differentiation. **(A–B)** Binned expression maps of *lamp1* mRNA abundance and *lamp1* mRNA association with polysomes across the germarium derived from scRNA-seq datasets. Dark blue indicates higher expression/association, and light blue indicates lower expression/association. **(C–D)** Confocal images of germaria expressing *UAS–lamp1::GFP* driven by *nos*GAL4. GFP fluorescence indicates Lamp1-positive vesicles, which are enriched during cyst stages. Scale bar, 25 µm. **(E–H)** Confocal images of an 8-cell cyst *from UAS–lamp1::GFP* germaria driven by *nos*GAL4, co-stained for Bruno to mark cyst stages and mAb414 to label NPCs. Yellow arrowheads indicate Lamp1-positive vesicles overlapping with NPC signal. Colocalization was assessed using Imaris 10.2 with intensity thresholds of 26 for mAb414 and 27 for GFP. Approximately 10.36% of Lamp1-positive vesicles overlapped with NPC signal. Scale bar, 5 µm. **(I–L)** Confocal images of an 8-cell cyst from *UAS–lamp1::GFP* germaria driven by *nos*GAL4, co-stained for Bruno and WGA to label NPCs. Yellow arrowheads indicate Lamp1-positive vesicles overlapping with NPC signal. Colocalization was assessed using Imaris 10.2 with intensity thresholds of 26 for WGA and 27 for GFP. Approximately 17.34% of Lamp1-positive vesicles overlapped with NPC signal. Scale bar, 5 µm.

**Figure 5S1:**
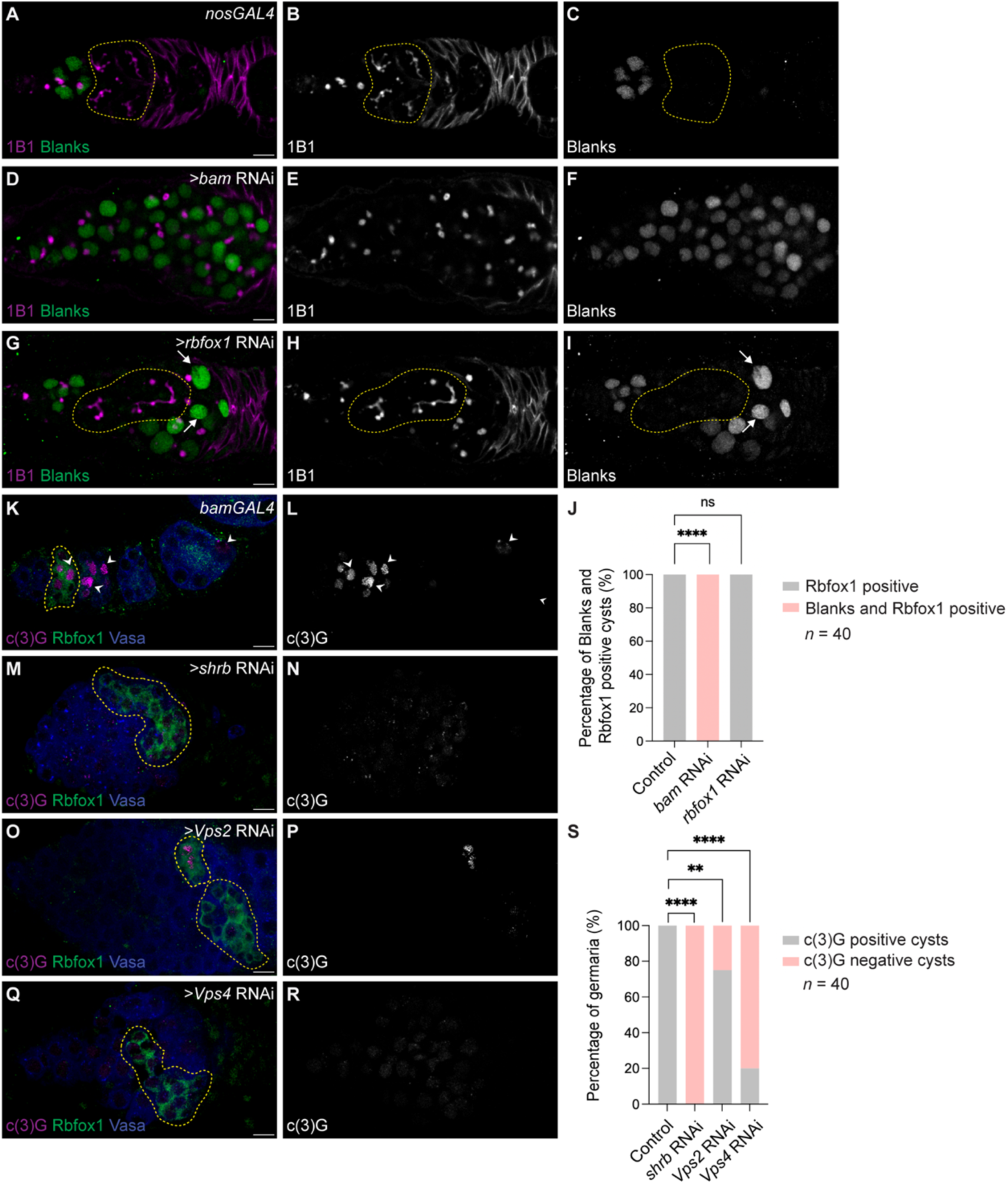
NPC degradation is required for initiation of meiotic recombination. **(A–I)** Confocal images of control, *bam*-depleted and *rbfox1*-depleted stained for Blanks, a germ cell gene marker. *rbfox1* depletion results in accumulation of cysts, whereas *bam* depletion leads to accumulation of GSCs and cystoblasts. White arrows indicate Blanks-positive single germ cells. Yellow dashed outlines mark 8-cell cysts that do not express Blanks. Scale bars, 10 µm. **(J)** Quantification of the percentage of germaria exhibiting extended Blanks expression in control, *bam*-, and *rbfox1*-depleted ovaries. Statistical analysis was performed using Fisher’s exact test. *n* = 40 germaria per genotype; ****, *p* < 0.0001. **(K–R)** Confocal images of control and germline-specific depletion of *shrb*, *Vps2*, or *Vps4* in germaria stained for crossover suppressor on 3 of Gowen [c(3)G], a marker of synaptonemal complex assembly and meiotic entry. c(3)G-positive cysts are observed in control germaria (white arrowheads) but are absent following depletion of *shrb*, *Vps2*, or *Vps4*. Yellow dashed outlines mark Rbfox1-positive 8-cell cysts. Scale bars, 10 µm. **(S)** Quantification of the percentage of germaria containing c(3)G-positive cysts in control and *shrb*-, *Vps*2-, or *Vps4*-depleted ovaries. Statistical analysis was performed using Fisher’s exact test. *n* = 40 germaria per genotype; ****, *p* < 0.0001; **, *p* = 0.0010.

**Fig. 6S1:**
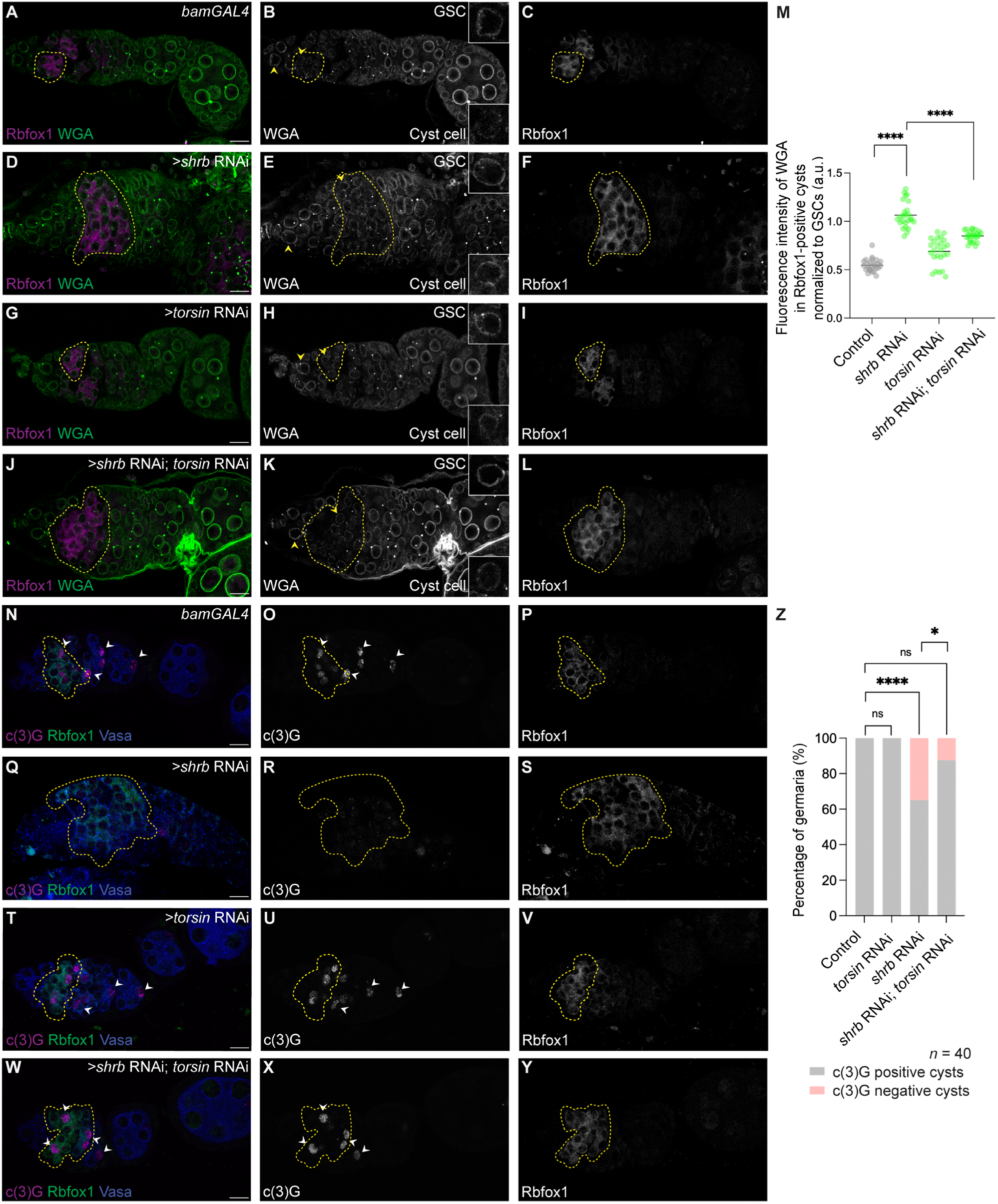
Forced NPC reduction during cyst differentiation rescues meiotic recombination defects caused by loss of ESCRT-III. **(A–I)** Confocal images of control, *shrb*-depleted, and *torsin*-depleted germaria stained with WGA to label NPCs and Rbfox1 to identify cyst stages. GSCs are indicated by yellow arrowheads. **(J–L)** Confocal images of *shrb* and *torsin* double-depleted germaria showing reduced NPC signal in cysts following forced NPC reduction during cyst differentiation. GSCs are indicated by yellow arrowheads. **(M)** Quantification of WGA fluorescence intensity in Rbfox1-positive cysts, normalized to GSCs within the same germarium and expressed in arbitrary units (a.u.). Statistical analysis was performed using a two-tailed unpaired Welch’s *t*-test. *n* = 5 germaria per genotype; ****, *p* < 0.0001. **(N–V)** Confocal images of control, *shrb*-depleted, and *torsin*-depleted germaria stained for c(3)G to assess initiation of meiotic recombination. White arrowheads indicate c(3)G-positive cells. **(W–Y)** Confocal images of germaria with combined depletion of *shrb* and *torsin* restores initiation of meiotic recombination in the *shrb*-depleted background. **(Z)** Quantification of the percentage of germaria containing c(3)G-positive cysts in control, *shrb*-, *torsin*-, and *shrb*/*torsin* double-depleted ovaries. Statistical analysis was performed using Fisher’s exact test. *n* = 40 germaria per genotype; ns, *p* > 0.9999; *, *p* = 0.0339; ****, *p* < 0.0001. Yellow dashed outlines mark Rbfox1-positive 8-cell cysts. Scale bars, 10 µm.

**Figure 6S2:**
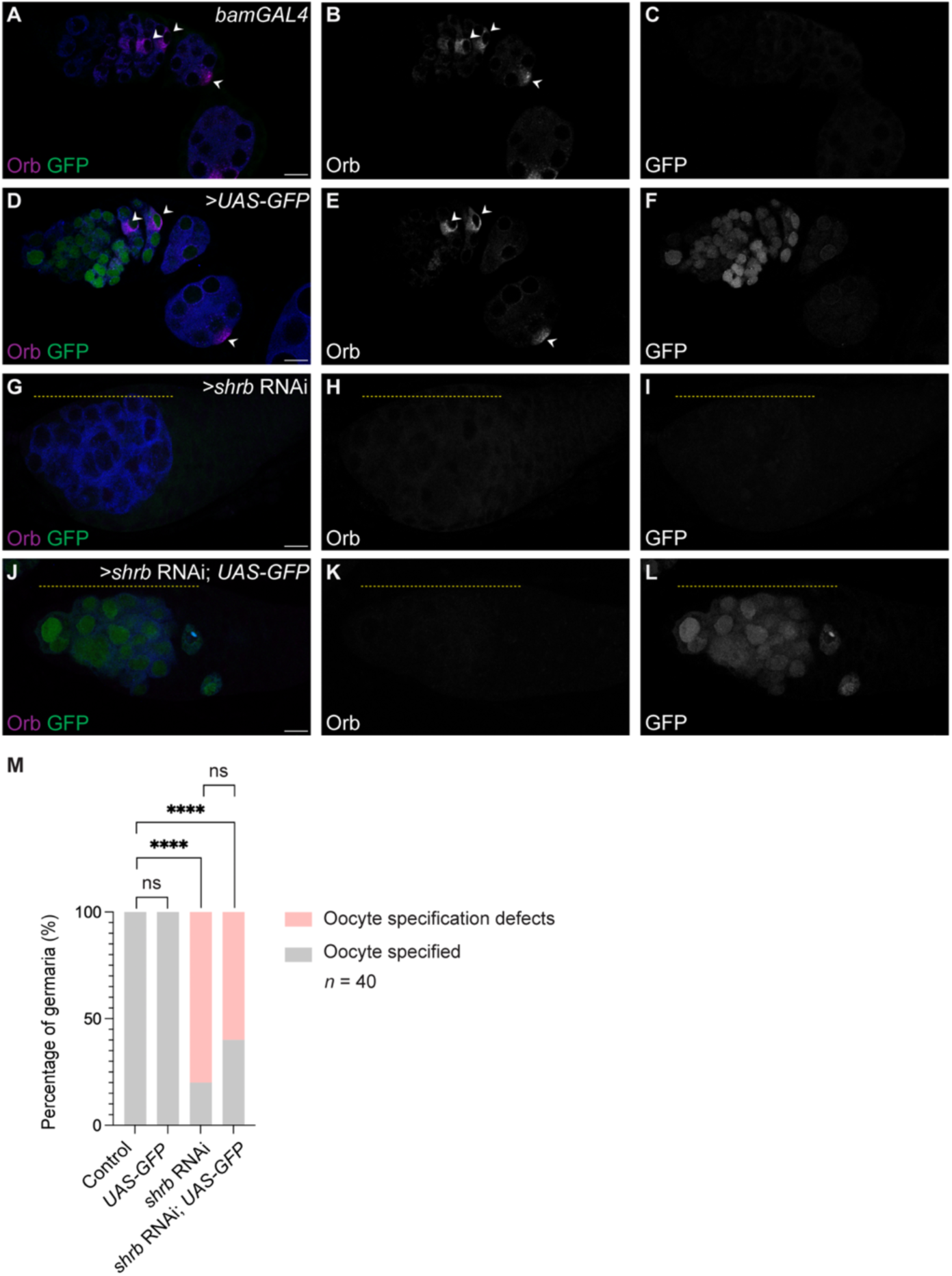
Addition of an extra UAS transgene does not rescue oocyte specification defects caused by loss of *shrb*. (A–I) Confocal images of control germaria, germaria expressing *UAS–GFP* under *bamGAL4* control, and *shrb*-depleted germaria, stained for Orb and GFP to assess oocyte specification. In *shrb*-depleted germaria, Orb fails to localize to a single cell within the cyst (yellow dashed outlines), indicating defective oocyte specification. In contrast, proper Orb localization to a single cell is observed in control germaria and in germaria expressing UAS–GFP alone (white arrowheads). (J–L) Confocal images of germaria expressing both *UAS–GFP* and *shrb* RNAi under *bamGAL4* control. Orb fails to localize to a single cell, indicating that the presence of an additional UAS transgene does not rescue the oocyte specification defect caused by *shrb* depletion. (M) Quantification of the percentage of germaria exhibiting a specified oocyte in control, *UAS–GFP*–expressing, *shrb*-depleted, and *UAS–GFP;shrb RNAi* germaria. Statistical analysis was performed using Fisher’s exact test. *n* = 40 germaria per genotype; ns, *p* > 0.9999; ****, *p* < 0.0001. Scale bars, 10 µm.

**Figure 6S3:**
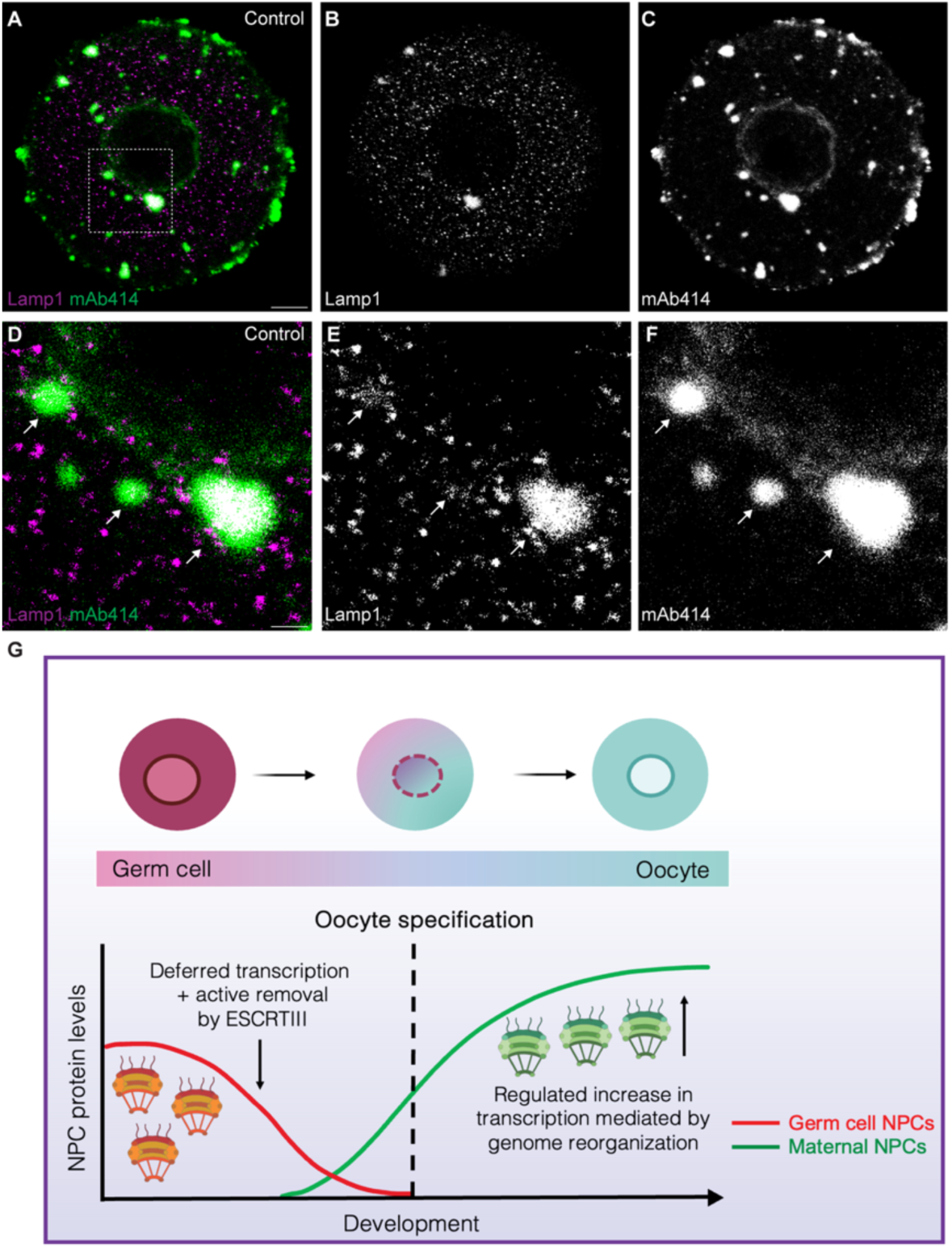
Nuclear pore complexes associate with lysosomes in prophase I–arrested mouse oocytes. **(A–C)** Confocal images of prophase I–arrested murine oocytes stained with mAb414 to label NPCs and Lamp1 to mark lysosomes. **(D–F)** Higher-magnification views highlighting regions of overlap between NPC and lysosome signals adjacent to the nuclear envelope (arrows). 100% overlap of puncta ∼4μm (n = 5 oocytes). Scale bars, 12.5 µm. **(G)** Model illustrating developmentally programmed replacement of germ cell NPCs with maternal NPCs during oocyte specification. Germ cell NPCs are reduced through a combination of passive dilution driven by deferred nucleoporin transcription and active removal from the nuclear envelope mediated by the ESCRT-III/Vps4 pathway. Subsequent upregulation of nucleoporin transcription supports assembly of maternal NPCs, which promote large-scale genome reorganization and establishment of the maternal chromatin state.

**Supplementary Table 1. RNA-seq and polysome-seq analysis of nucleoporin (Nup) transcripts.** RNA-seq and polysome profiling–coupled RNA sequencing (polysome-seq) data for 29 nucleoporin genes analyzed in ovaries.

## Notes

### Competing Interest Statement

The authors have declared no competing interest.

